# Epithelial density controls cell migration through an adhesion-nucleus mechanotransduction pathway

**DOI:** 10.64898/2026.04.22.720102

**Authors:** Louis Laurent, Thomas Germier, Nicolas Audugé, Helena Canever, Joséphine Schelle, Hugo Lachuer, Philippe P. Girard, Jean-Baptiste Manneville, François Sipieter, Nicolas Borghi

## Abstract

Cell density is thought to regulate tissue growth, homeostasis, and regeneration, yet how cells sense and respond to density remains poorly understood. To investigate density-dependent mechanotransduction, we combined genetically encoded biosensors with quantitative fluorescence microscopy in epithelial cells subjected to genetic, pharmacological, and mechanical perturbations. We found that low epithelial density promotes focal adhesion growth, causing mechanical relaxation of vinculin and release of its competitive binding with FAK and ERK. This enables FAK to bind and activate ERK in the cytoplasm. Cytoskeletal tension transmitted through LINC complexes drives their nuclear translocation, where low density induces ERK-dependent chromatin decondensation and increased nuclear envelope tension. This recruits and activates cPLA2, leading to arachidonic acid production and enhanced cell migration. Together, these findings identify a mechanochemical pathway linking cell density to epithelial migration via ERK, cPLA2, and mechanically regulated signaling from adhesions to the nucleus.

**Significance statement:** In multicellular organisms, cells constantly experience crowding, yet how they detect changes in cell density and convert them into biological responses is not well understood. Here, we show how low cell density mechanically triggers signaling inside epithelial cells to promote migration. When cells are sparse, adhesion sites grow and release key signaling proteins that move into the nucleus stretched by the cytoskeleton. These mechanical and biochemical inputs alter nuclear structure, activates lipid signaling, and boosts cell migration. By revealing how mechanical forces, cell adhesions, and nuclear signaling work together, this study provides a clear mechanistic link between cell density and migration, a process central to tissue growth, repair, and disease progression.

## Introduction

Simple epithelia are probably the most ancestral and archetypal tissues of multicellular organisms (1). The perception of cell density within epithelia is a fundamental property governing tissue growth, homeostasis and repair (2). While high epithelial cell density induces quiescence and arrest in cell motion (3, 4), low cell density promotes cell proliferation and migration (5), thereby leading to an optimal density at which epithelial functions operate. The Extracellular-signal Regulated Kinase (ERK), which serves as a major hub for the regulation of cell behavior, senses epithelial density and regulates cell migration accordingly (6). However, the mechanisms by which it does so remain largely unexplored.

Mechanical stresses inherent in changes in cell density and migration make mechanotransducers ideal candidates for the regulation of ERK. Consistently, intercellular adhesion proteins E-cadherins and ion channels Piezo1 have been involved in the mechanical induction of ERK activity by receptor tyrosine kinases in various models (7). Proteins of Focal Adhesions (FAs), which enable cells to adhere to the extra-cellular matrix, are also long known regulators of ERK activity (8). Vinculin was notably found to control Focal Adhesion Kinase (FAK)-Paxillin interactions and subsequent ERK activity upstream of cell migration in fibroblasts (9). While vinculin ensures force transmission between the cytoskeleton and the matrix (10), it is however unknown whether vinculin mechanosensitivity is involved in the regulation of ERK activity or its dependence on cell density.

The regulation of cell migration by ERK as a response to epithelial cell density ultimately involves actomyosin contractility (6, 11). While ERK might do so through direct phosphorylation of myosin light chain kinase as in other cell types (12), a major consequence of ERK activation is its translocation from the cytoplasm into the nucleus (13). Mechanical regulation of nucleo-cytoplasmic shuttling is a critical step in many signaling pathways (14, 15), and the nucleus is now recognized as a mechanosensor capable of regulating amoeboid migration through the activation of the phospholipase cPLA2 (16, 17). Interstingly, cPLA2 is also a target of ERK (18), and its deregulation promotes epithelium-to-mesenchyme transition in carcinomas (19). This opens the possibility that the regulation of epithelial cell migration by a mechanostransuction pathway involving ERK and cPLA2 is a physiolpgical response to cell density that is deregulated in disease.

Here, we sought to assess whether FAs could serve as a cell density sensor upstream of ERK activity in epithelia, and whether epithelial migration could in turn rely on nuclear mechnaotransduction pathways dependent on ERK. To test this, we combined various fluorescent biosensors of mechanical and biochemical activities and quantitative fluorescent imaging approaches, together with specific pharmacological, genetic and mechanical perturbations in a model of simple epithelia in culture. Our results support a model in which low epithelial cell density triggers the release of active ERK in the cytoplasm, promotes its translocation into the nucleus, which results in cPLA2 activation, upstream of epithelial cell migration.

## Results

### The dependence of ERK on epithelial cell density relies on Focal adhesion growth and vinculin tension

To assess whether vinculin is involved in the sensitivity of ERK activity to epithelial density, we plated Wild-Type (WT) and vinculin-KO MDCK cells expressing a sensor of ERK activity (ERK-KTR, see Materials and Methods) at various levels of confluence between 1000 and 3000 cells*/*mm^2^ (Fig. 1A, Fig. S1A, B). WT cells exhibited a significant cytoplasmic translocation of ERK-KTR from high to low density, indicative of an increase in ERK activation, while KO cells exhibited an intermediate state independent of cell density that was higher than in WT cells at high density. Remarkably, the transition in ERK activity in WT cells was rather abrupt around 1600 cells*/*mm^2^. Furthermore, re-expressing a vinculin construct bearing a tension sensor (VinTS) in KO cells rescued the lower ERK activity to similar levels as in WT cells (Fig. 1C). These results support that vinculin controls the dependence of ERK activity on epithelial cell density.

**Fig. 1.**
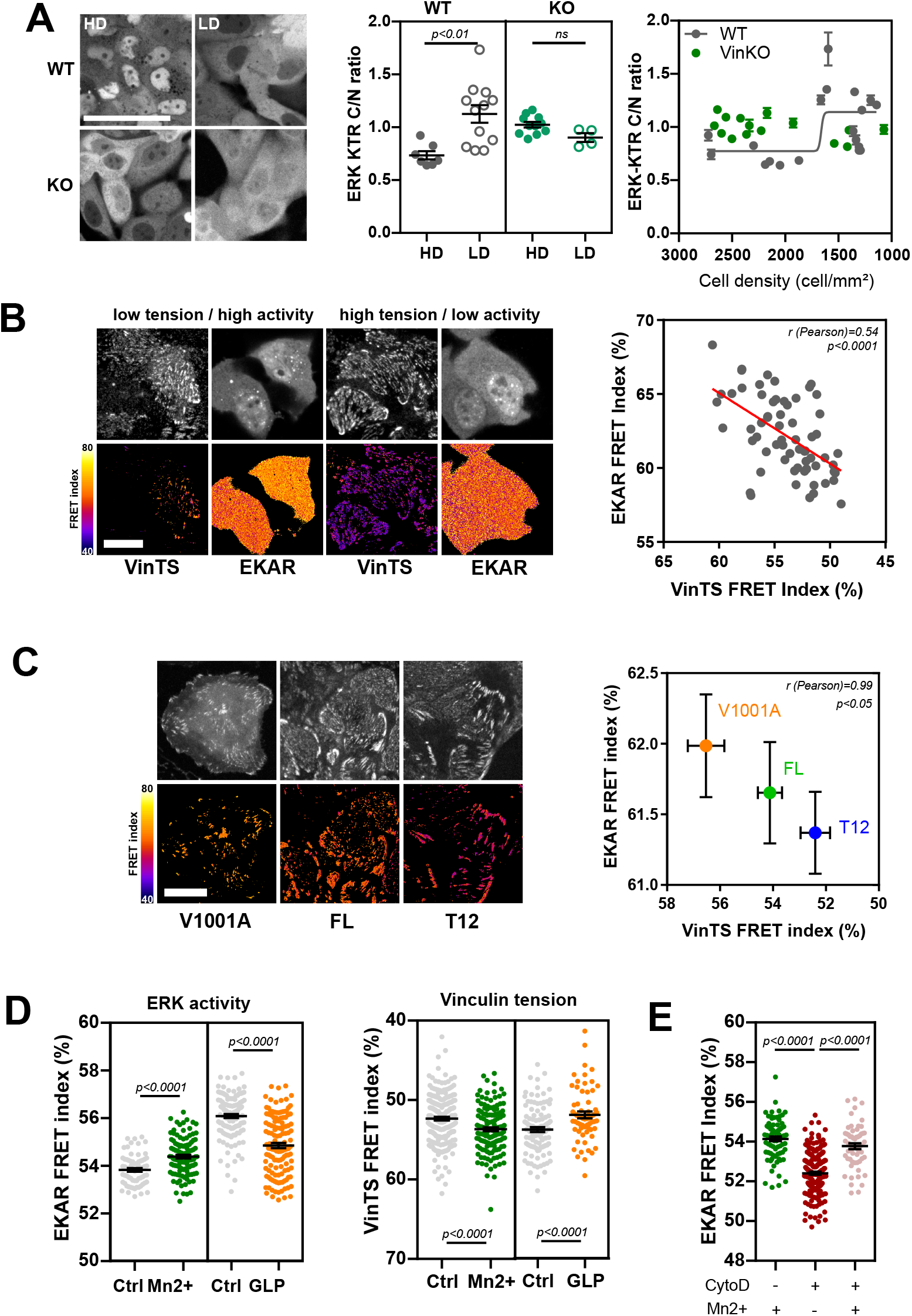
(A) Left: fluorescence image of MDCK cells expressing the ERK KTR sensor, WT: wild-type, KO: Vinculin KO. Scale bar = 50*µ*m. Right: ERK KTR cytoplasm/nucleus (C/N) intensity ratio as a function of epithelial cell density (HD: high density >1600 cells/mm^2^, LD: low density <1600 cells/mm^2^). Mean*±*SEM of 19 (WT) and 15 (KO) fields of view (FOV) from 2 (WT) and 3 (KO) independent experiments. A FOV contains about 150-200 cells at HD and twice less at LD. The curve is a sigmoidal fit. (B) Left: fluorescence and FRET images of MDCK cells expressing spectrally compatible VinTS (TFP-Venus) and EKAR (Ruby-Kate) sensors. Scale bar = 20*µ*m. Right: spontaneous variability of EKAR and VinTS FRET indices within a cell population. 64 cells from 3 independent experiments. (C) Right: fluorescence and FRET images of MDCK cells expressing the original VinTS (FL) and the V1001A and T12 mutants. Scale bar = 20*µ*m. Right: FRET indices of EKAR and VinTS sensors as a function of vinculin mutants. Mean±SEM of 66 (FL), 75 (T12) and 45 (V1001A) cells from 3 independent experiments. (D) EKAR (left) and VinTS (right) FRET indices as a function of integrin activation (Mn^2+^) or inhibition (GLP). Mean SEM of 70 (Ctrl/Mn^2+^), 110 (Mn^2+^), 46 (Ctrl/GLP), 48 (GLP) cells from 3 independent experiments (EKAR). 201 (Ctrl/Mn^2+^), 138 (Mn^2+^), 104 (Ctrl/GLP), 51 (GLP) cells from 3 independent experiments (VinTS). (E) EKAR FRET index as a function of integrin activation (Mn^2+^) and/or cytoskeleton disruption (CytoD). Mean*±*SEM of 77 (Mn^2+^), 211 (CytoD) and 57 (CytoD+Mn^2+^) cells from 3 (Mn^2+^) or 2 independent experiments. Two-tailed Mann-Whitney tests (A, D, E). Pearson correlation (B, C).

Because vinculin is a mechanosensor (10), we then tested whether its tension relates to ERK activity. For this, we simultaneously assessed vinculin tension and ERK activity in cells that expressed VinTS together with a spectrally compatible FRET sensor of ERK activity (EKAR, see Materials and Methods) bearing red and far-red fluorophores (Fig. S1D). We found that cells spontaneously exhibited a higher ERK activity with a lower vinculin tension (Fig. 1B). To test whether changes in vinculin tension can cause changes in ERK activity, we expressed mutants versions of VinTS that impaired (V1001A) or promoted (T12) the interaction with F-actin (Fig. S1E), together with the EKAR sensor. We found that expression of the relaxed, V1001A mutant caused an increase in ERK activity while the tensed, T12 mutant decreased ERK activity, compared to the control VinTS (FL) (Fig. 1C). These results support that vinculin tension relaxation promotes ERK activation.

To further assess how epithelial cell density can control ERK activity through vinculin tension, we tested whether the growth of Focal Adhesions (FAs) was involved. To this end, we activated integrin interaction with the extracellular matrix with Mn^2+^, which increased the size of FAs, as low epithelial cell density did (Fig. S1F, G). Alternatively, we inhibited integrin interaction with the extracellular matrix with GLPG0187 (GLP), which decreased the size of FAs, as high epithelial cell density did. In cells expressing VinTS and EKAR, we found that activating integrins decreased vinculin tension and, as expected, increased ERK activity (Fig. 1D). Consistently, inhibiting integrins increased vinculin tension and decreased ERK activity (Fig. 1D). Together, these results support that low density promotes the growth of relaxed FAs, which in turns activates ERK. Finally, we reasoned that if cytoskeletal tension is dispensable for ERK activation upon FA growth, it would be possible to activate ERK by activation of integrins in the absence of an intact cytoskeleton. Indeed, ERK activity reached similar levels upon integrin activation whether or not cells were inhibited for actin polymerisation by cytochalasin D (CytoD) (Fig. 1E).

### Low density dissociates ERK from Focal Adhesions and promotes the formation of an active ERK/FAK complex in the cytoplasm

Because interactions between ERK and FA proteins regulate ERK activity (9), we wondered whether cell density, through its effect on vinculin tension, could modulate these interactions. To test this, we assessed direct interactions between fluorescent constructs of vinculin (VCL), paxillin (PXL), Focal Adhesion Kinase (FAK) and ERK by FLIM-FRET as a function of cell density (Fig. 2A). We found that in FAs, while vinculin and paxillin interacted with the other three proteins at high density, paxillin only interacted with vinculin and FAK, and vinculin with paxillin, at low density (Fig 2B, S2A,B). Moreover, FAK did not interact directly with ERK in FAs, regardless of cell density (Fig. S2C). In the cytoplasm, vinculin, paxillin and FAK had the same interacting partners as in FAs at high density (Fig. 2C, S2D, E). At low density, however, ERK and FAK now interacted with each other directly, and each with paxillin, while vinculin interacted with none of them (Fig. 2C, S2D, E). Finally, because FAK activity promotes ERK activity (9), we tested whether it was required for the dependence of ERK activity on density. As expected, pharmacological inhibition of FAK abolished ERK activity and its dependence on cell density (Fig. 2D).

**Fig. 2.**
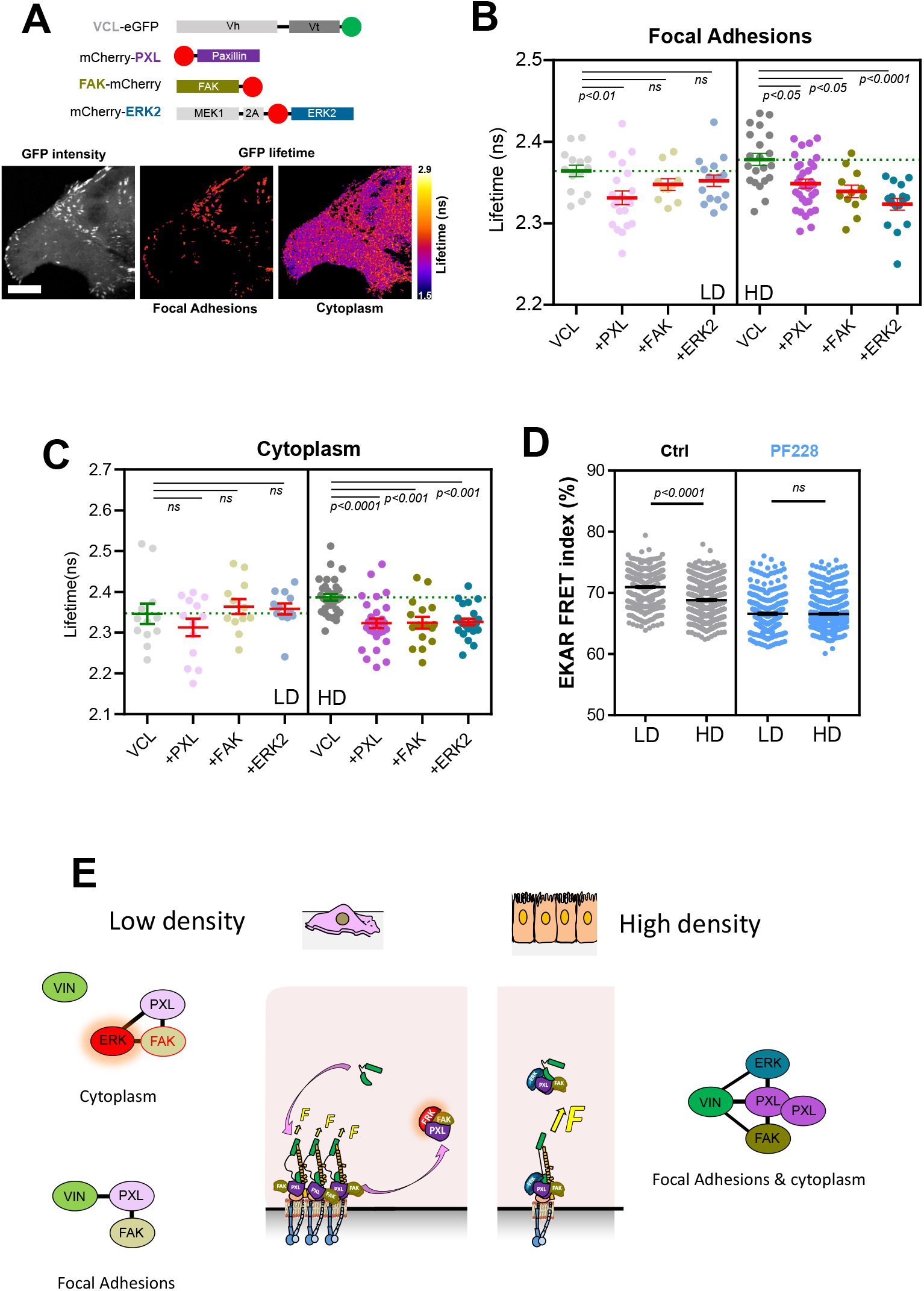
(A) Top: sketch of the constructs used for FRET-FLIM experiments. Bottom: typical fluorescence intensity and FLIM images of a cell expressing VCL-eGFP. Scale bar = 10*µ*m. (B) Fluorescence lifetime of VCL-eGFP in Focal Adhesions of cells expressing another mCherry construct or none at low (LD) and high (HD) cell density. Mean*±*SEM of 14 (LD, VCL), 21 (LD, +PXL), 10 (LD, +FAK), 16 (LD, +ERK), 21 (HD, VCL), 32 (HD, +PXL), 12 (HD, +FAK) and 16 (HD, +ERK) cells (*>* 50 FAs) from 3 independent experiments. (C) Fluorescence lifetime of VCL-eGFP in the cytoplasm of cells expressing another mCherry construct or none at low (LD) and high (HD) cell density. Mean*±*SEM of 12 (LD, VCL), 13 (LD, +PXL), 12 (LD, +FAK), 12 (LD, +ERK), 31 (HD, VCL), 27 (HD, +PXL), 16 (HD, +FAK), 24 (HD, +ERK) cells from 3 independent experiments. (D) EKAR FRET index at low and high cell density exposed or not to the FAK inhibitor PF228. Mean*±*SEM of *>* 300 cells from 3 independent experiments. (E) Working model of direct interactions in the cytoplasm and Focal Adhesions as a function of cell density. Kruskal-Wallis and Dunn’s multiple comparisons tests (B, C), two-tailed Mann-Whitney tests (D).

Altogether, these results are consistent with a model by which low density alters force-sensitive interactions in FAs in a way that allows the release of a cytoplasmic complex in which a direct interaction between active FAK and ERK activates ERK (Fig. 2E).

### Focal adhesion growth promotes ERK and FAK nuclear translocation in a manner dependent on the cytoskeleton and nesprin tension

Nuclear translocation is a typical outcome of ERK activation in epithelial tissues (6). Therefore, we sought to test whether it depended on epithelial density through FA growth. Using an ERK2Loc construct (20), we confirmed that ERK localization was more nuclear at low cell density than at high density (Fig. 3A). Moreover, activation of integrins (Mn^2+^) also increased ERK nuclear localization, while their inhibition (GLP) did not (Fig. 3B). The nuclear translocation of ERK upon FA growth reached a plateau within 10 min, a similar time scale as if ERK was activated by the protein kinase C (PKC) with Phorbol 12-Myristate 13-Acetate (PMA) (Fig. S3A). The use of a FRET sensor of ERK activity targeted to the nucleus (EKAR-NLS) showed that nuclear ERK activity followed the same kinetics, whether upon integrin or PKC activation (Fig. S3B).

**Fig. 3.**
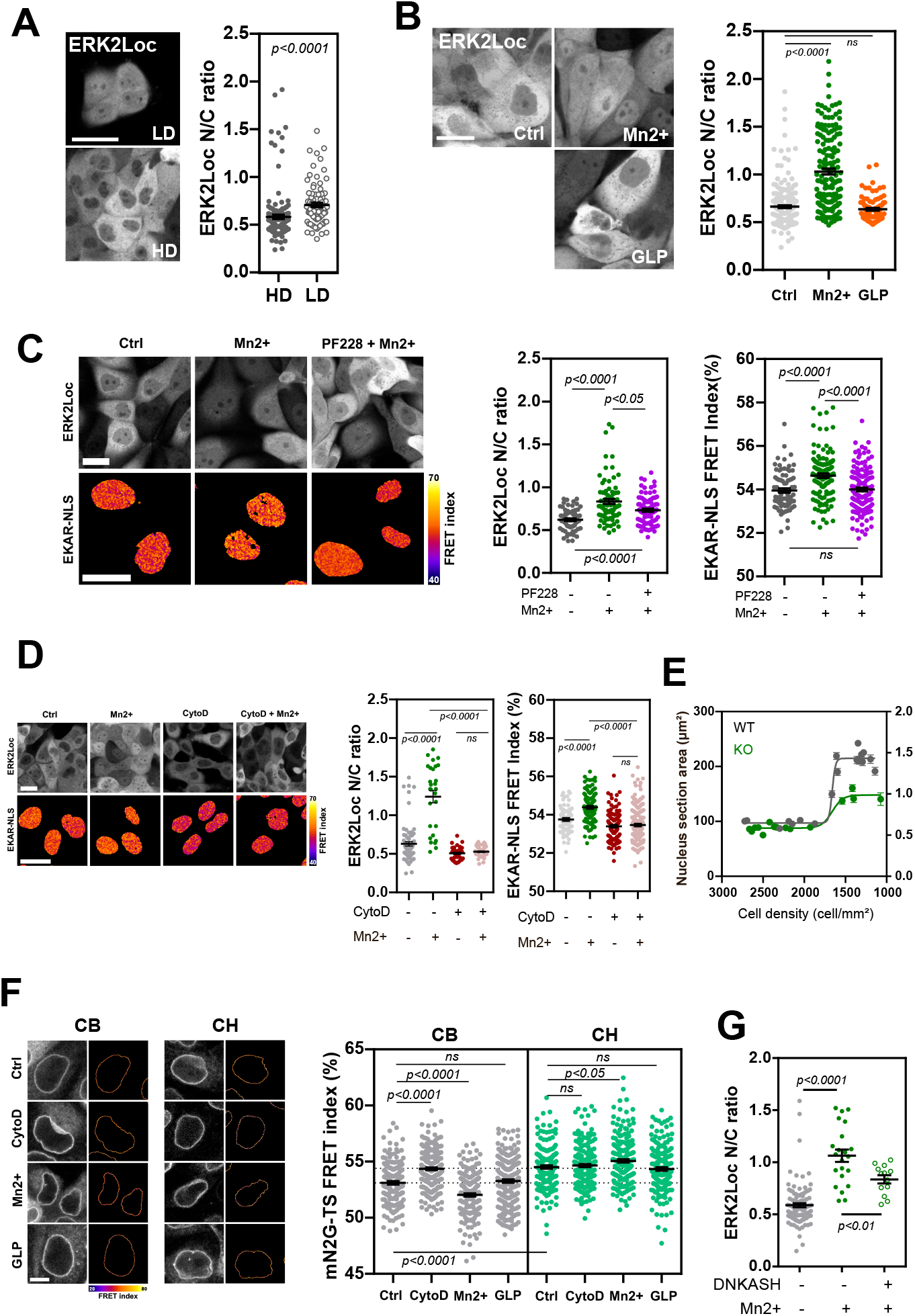
(A) Left: fluorescence images of MDCK cells expressing the ERK2Loc construct as a function of cell density. Scale bar = 50*µ*m. Right: ERK2Loc nucleus/cytoplasm (N/C) intensity ratio (HD: high density, LD: low density). Mean±SEM of 153 (HD) and 88 (LD) cells from 2 independent experiments. (B) Left: Fluorescence images of MDCK cells expressing the ERK2Loc construct as a function of integrin activation (Mn^2+^) and inhibition (GLP). Scale bar = 20*µ*m. Right: ERK2Loc nucleus/cytoplasm (N/C) intensity ratio. Mean±SEM of 235 (Ctrl), 166 (Mn^2+^) and 93 (GLP) cells from 3 independent experiments. (C) Left: fluorescence images of cells expressing ERK2Loc and FRET images of cells expressing EKAR-NLS as a function of integrin activation (Mn^2+^) and/or FAK inhibition (PF228). Scale bar = 20*µ*m. ERK2Loc N/C ratio (middle) and EKAR-NLS FRET index (right) in these conditions. Mean ±SEM of 67 (Ctrl), 71 (Mn^2+^) and 87 (PF228+Mn^2+^) cells from 3 independent experiments (ERK2Loc), and 91 (Ctrl), 121 (Mn^2+^) and 134 (PF228+Mn^2+^) cells from 3 independent experiments (EKAR-NLS). (D) Left: fluorescence images of cells expressing ERK2Loc and FRET images of cells expressing EKAR-NLS as a function of integrin activation (Mn^2+^) and/or cytoskeleton disruption (CytoD). Scale bar = 20*µ*m. ERK2Loc N/C ratio (middle) and EKAR-NLS FRET index (right) in these conditions. Mean± SEM of 76 (Ctrl), 28 (Mn^2+^), 35 (CytoD) and 41 (CytoD+Mn^2+^) cells from 2 independent experiments (ERK2Loc), and 79 (Ctrl), 197 (Mn^2+^), 110 (CytoD) and 171 (CytoD+Mn^2+^) cells from 3 (Ctrl, Mn^2+^) or 2 (CytoD, CytoD+Mn^2+^) independent experiments (EKAR-NLS). (E) Nucleus section area as a function of epithelial cell density in WT and KO cells. Mean*±*SEM of 19 (WT) and 15 (KO) FOVs from 2 (WT) and 3 (KO) independent experiments (same as in Fig. 1A), Curves are sigmoidal fits. (F) Left: Fluorescence and FRET images of cells expressing the mNesprin-TSMod (mN2G-TS) cytoskeleton-binding (CB) and calponin homology domain (CH) constructs as a function of cytoskeleton disruption (CytoD) or of integrin activation (Mn^2+^) or inhibition (GLP). Scale bar = 10*µ*m. Right: FRET indices of the mN2G-TS CB and CH constructs in these conditions. Mean ± SEM of 154 (Ctrl), 203 (CytoD), 181 (Mn^2+^) and 212 (GLP) cells from 3 independent experiments (CB), and 179 (Ctrl), 195 (CytoD), 203 (Mn^2+^) and 190 (GLP) cells from 3 independent experiments (CH). Dotted lines show the Ctrl/CB and Ctrl/CH levels to facilitate comparisons with other conditions. (G) ERK2Loc N/C ratio in cells expressing or not the nesprin dominant negative construct DNKASH with or without activation of integrins (Mn^2+^). Mean*±*SEM of 106 (Ctrl), 22 (Mn^2+^) and 12 (DNKASH+Mn^2+^) cells from 2 (Ctrl) or 1 independent experiments. Two-tailed Mann-Whitney tests.

Next, we assessed the dependence of nuclear localization and activity of ERK on FAK, which was required for its cytoplasmic activity (see Fig.2D). As expected, inhibition of FAK in cells exposed to integrin activation reverted ERK nuclear localization and activity to lower levels, as in cells the integrins of which were not activated (Fig. 3C). Interestingly, in cells expressing FAK-GFP and ERK2Loc, we observed a correlation in their nucleo-cytoplasmic translocation in cells that were exposed to integrin activation, inhibition, or left at rest, while cells activated through PKC exhibited no such correlation (Fig. S3C). This supports that activation of ERK by FAs results in a specific response at the nuclear level by bringing FAK along.

The nuclear translocation of a number of master regulators of cell fate and behavior is a mechanosensitive process though to involve the dilation of nuclear pores as a result of extra-cellular or cytoskeletal forces deforming the nucleus, in part through LINC complex (14, 15). To assess how this applies to ERK, we first assessed how its translocation depended on the integrity of the cytoskeleton. Consistently, CytoD treatment abolished the nuclear translocation and activity of ERK induced by integrin activation and brought them lower than their basal levels in less that 10 min (Fig. 3D, S3D). Then, we examined nuclear deformations and found that the nucleus cross-section area increased as epithelial cell density decreased, with a sharp transition around 1600 cells*/*mm^2^, an effect largely impaired in cells lacking vinculin (Fig. 3E). Altogether, these results support a model by which FA growth, in addition to activating ERK at low cell density (see Fig. 1), enables cytoskeleton-dependent nuclear deformations that facilitate the nuclear translocation of ERK.

The active transport of ERK into the nucleus primarily relies on importin-7, but may also involve importin-*β* (21). Consistently, we found that the nucleo-cytoplasmic ratio of importin-7 correlated with the nucleus section area (Fig. S3E) and increased in cells exposed to integrin (Mn^2+^) or PKC (PMA) activation (Fig. S3F). Moreover, exposure of cells to ivermectin, an inhibitor of importin*−α/β* (22), partially reduced ERK nuclear translocation when induced by PKC while it did not when induced by FA growth (Fig. S3G). These results support a selective use of importins depending on the level or the mechanism of ERK activation, with importin-*β* being dispensable for ERK translocation upon activation by FAs, but involved when ERK is activated by other pathways.

Finally, we though to assess whether cytoskeletal tension on the LINC complex could contribute to ERK nuclear translocation. Consistently, using a nesprin tension sensor (mN2GTS) (Fig. S3H), we found that nesprin tension was dependent on the integrity of the cytoskeleton and was increased upon FA growth (Mn^2+^) (Fig. 3F). Moreover, expression of a dominant negative nesprin (DNKASH, Fig. S3H) that disrupts interactions between LINC complexes and the cytoskeleton (23), decreased the nuclear translocation of ERK induced by FA growth (Fig. 3G), and by PKC (Fig. S3I). Altogether, these results support that cytoskeletal tension exerted on LINC complexes upon FA growth facilitates the nuclear translocation of ERK, regardless of the activating signal.

### Low cell density induces ERK-dependent chromatin decondensation and increased nuclear envelope tension

ERK has many nuclear targets and its activity generally results in increased transcription. This associates with chromatin remodelling in a decondensed state by means of post-translational modifications (24). Thus, we thought to assess the effect of cell density on chromatin condensation and its dependence on ERK. To do so, we monitored the fluorescence lifetime of H4-GFP, which is sensitive to chromatin condensation (25). We found that chromatin was less condensed at low cell density and that the effect depended on ERK (Fig. 4A).

**Fig. 4.**
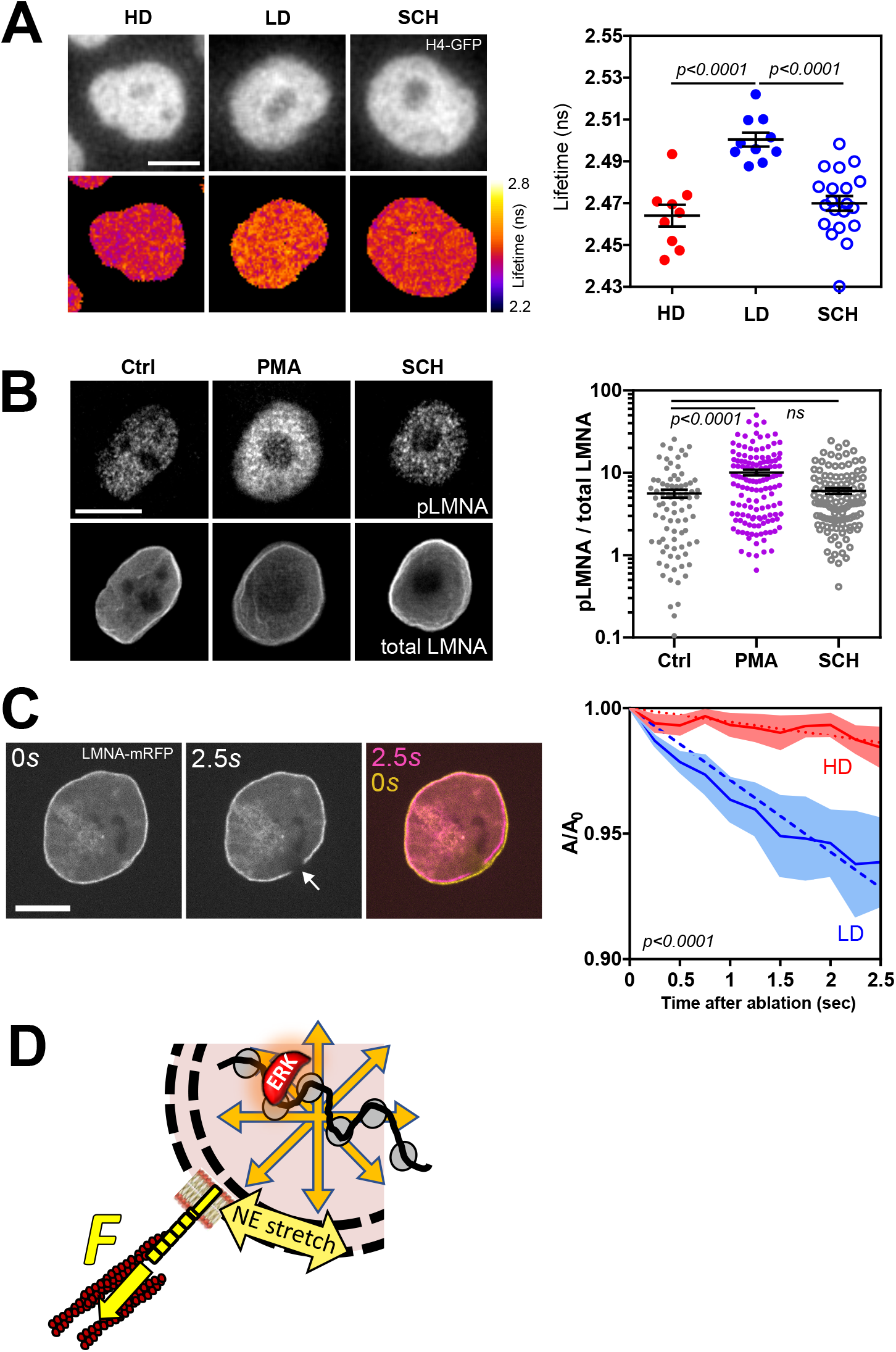
(A) Left: Images of fluorescence intensity and lifetime of H4-GFP in MDCK cells. Mean fluorescence lifetime of H4-GFP in cells at high (HD) and low (LD) density without or with inhibition of ERK (SCH). Mean*±*SEM of 9 (HD), 10 (LD) and 20 (SCH) cells from 1 (HD, LD), or 2 (SCH) experiments. (B) Left: Immonofluorescence images of phospho-Lamin A (pLMNA) and total Lamin A (total LMNA) in control, ERK activated (PMA), and ERK inhibited (SCH) cells. Right: intensity ratio (pLMNA / total LMNA) in those conditions. Mean*±*SEM of 83 (Ctrl), 135 (PMA) and 115 (SCH) cells from 2 experiments. (C) Left: LaminA-mCherry nucleus before and after ablation (arrow) of the NE (and overlay). Right: rate of nucleus deflation after laser ablation of the NE at low (LD) and high (HD) epithelial density. Mean*±*SEM of 13 (LD) and 21 (HD) ablations from 2 independent experiments. Dotted lines are lienar fits. (D) Working model of increased NE tension at low density (see text for details). Two-tailed Mann-Whitney (A, B) and extra sum-of-squares F tests (C). Scale bars = 10*µ*m.

A number of mechanisms participate to such chromatin remodelling. Recently, Ser22-phosphorylated lamin A/C have been implicated, by binding to enhancers near active genes, in regions rich in acetylated histone, a mark of decondensed chromatin (26). Interestingly, ERK phosphorylates Lamin A/C at Ser22 (27). We thought to test whether this was the case in MDCK cells by immunofluorescence of Ser22-phosphorylated laminA/C (pLMNA). Consistently, activation of ERK (PMA) increased the pLMNA/LMNA ratio (Fig. 4B). Moreover, pLMNA exhibited a pronounced localization at the nuclear interior rather than at the periphery, consistent with its association with active genes. Additionally, Ser22-phosphorylation of lamins promotes their mobility from the nuclear lamina (28). ERK activity should therefore do the same. Consistently, FRAP experiments showed increased fluorescence recovery of LMNA-mRFP at the nuclear lamina under ERK activation (PMA) compared to control or ERK inhibition (SCH), consistent with pLMNA being released from the nuclear lamina (Fig. S4A). Altogether, these results suggest that ERK promotes the release of pLMNA from the nuclear lamina to the nucleus interior, thereby providing a mechanism to remodel the chromatin.

Finally, we sought to assess whether chromatin decondensation at low density could increase the internal pressure on the nuclear envelope (NE), thereby contributing to increasing its tension (29). To test this, we performed localized laser ablation of the NE, in order to release the internal pressure without substantially affecting cytoskeletal tension through LINC complexes. The nucleus deflation rate immediately after ablation is proportional to the balance between the pressure-driven tension and the viscosity of the intranuclear fluid (30). We found that the nucleus deflation rate was about an order of magnitude larger in cells at low density (Fig. 4C). This is consistent with increased tension, provided chromatin decondensation did not also decreased the viscosity. To confirm this, we assessed the relaxation of nucleus indentations with intracellular optical tweezers (31) (Fig. S4B, C), which depends on the balance between nucleus stiffness and viscosity. We found that decondensing chromatin by inhibition of histone deacetylases (Sodium Butyrate) or by activation of ERK (PMA) did not affect nucleus viscosity (Fig. S4D).

Altogether, these results support that low density promotes the decondensation of chromatin in an ERK-dependent manner, which contributes to an increase in NE tension, in addition to LINC-mediated cytoskeletal forces (Fig. 4D).

### Low density induces ERK-dependent cPLA2 activity, which regulates epithelial cell migration

Deformation of the nucleus is known to trigger the recruitment of the phospholipase cPLA2 to tensed NE (32). We reasoned that the effects of density on nucleus shape and NE tension (see Fig. 3E, 4C) could impact cPLA2 localization too. Consistently, we found that cPLA2 was enriched at the NE at low density (Fig. 5A). Moreover, cPLA2 is also a substrate of ERK, which targets its serine 505 (33). Thus, we thought to assess the implication of ERK in cPLA2 activation. Using immunofluorescence, we found that the nuclear amount of pSer505-cPLA2 correlated with that of ERK, although not all ERK positive nuclei were positive for pSer505cPLA2 (Fig. 5B). Furthermore, cells plated at low density exhibited higher nuclear levels of pSer505-cPLA2 than cells at high density, and forcing the activation of ERK through PKC in cells at high density increased pSer505-cPLA2 to levels comparable to that in cells at low density (Fig. 5C). Altogether, these results support that ERK activation at low cell density induces cPLA2 activity through both phosphorylation and NE tension increase.

**Fig. 5.**
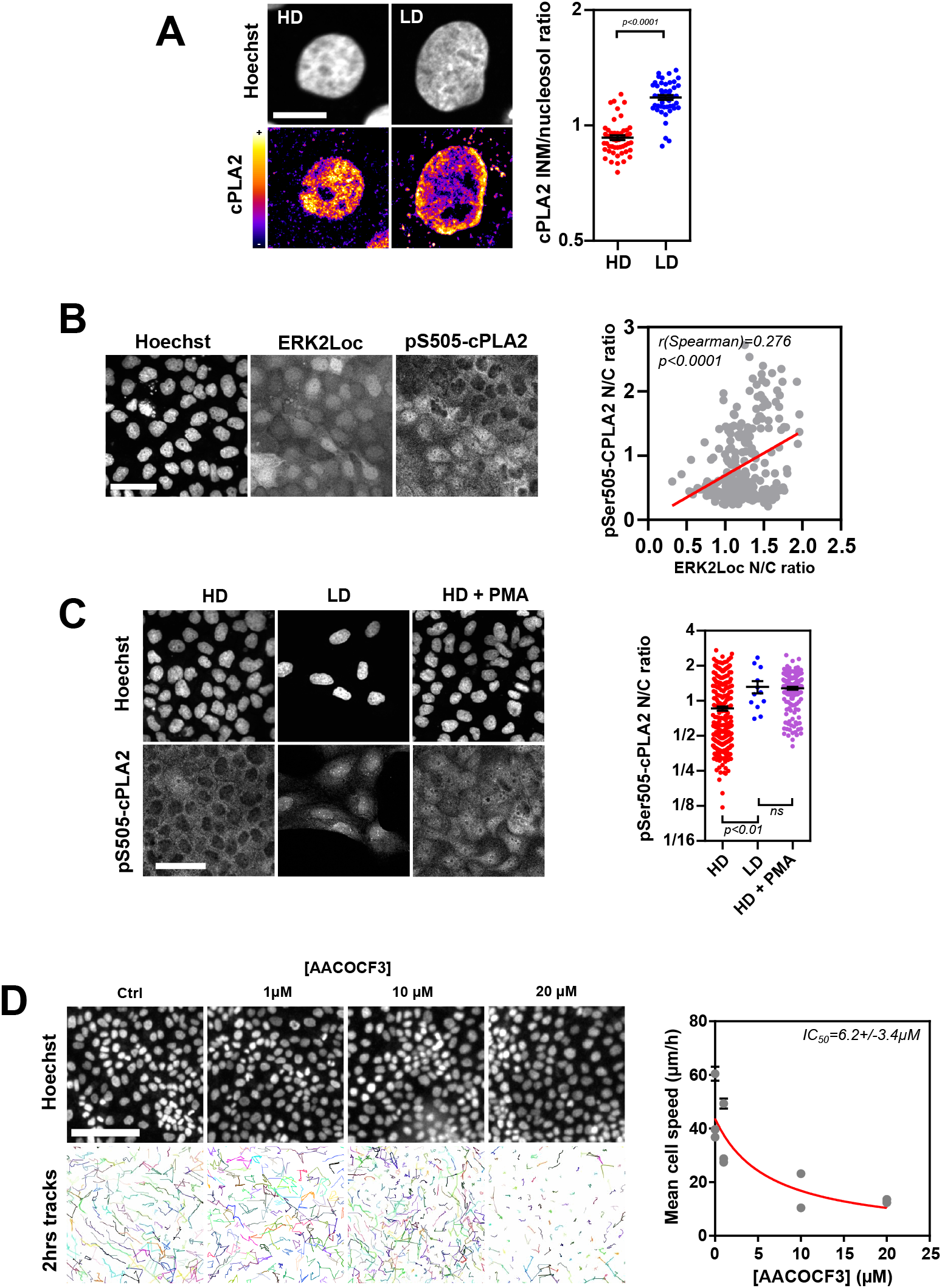
(A) Left: Immunofluorescence of cPLA2 in cells at high (HD) and low (LD) epithelial density. Scale bar = 10*µ*m. Right: Inner Nuclear Membrane (INM)/Nucleosol ratio of cPLA2 immunofluorescence intensity as a function of epithelial density. Mean*±*SEM of 52 (HD) and 49 (LD) cells from 2 experiments. (B) Left: Immunofluorescence of pS505-cPLA2 and fluorescence of ERK2Loc in confluent cells. Scale bar = 50*µ*m. Right: correlation of pS505-CPLA2 and ERK2Loc N/C ratios. 247 cells from 1 experiment. The straight line is a linear fit. (C) Left: Immunofluorescence of pS505-cPLA2 as a function of epithelial density and activation of ERK (PMA). Scale bar = 50*µ*m. Right: pS505-CPLA2 N/C ratios in these conditions. Mean*±*SEM of 259 (HD), 12 (LD) and 150 (HD+PMA) cells from 1 experiment. (D) Left: Fluorescence images of nucleus-stained cells at confluence under increasing amounts of cPLA2 inhibitor AACOCF3 and their corresponding migration tracks over 2 hours. Scale bar = 100*µ*m. Right: Mean cell speed of confluent cells over 2hrs. 1 point per FOV from 3 (Ctrl, 1*µ*M) and 2 (10, 20*µ*M) independent experiments, mean*±*SEM of 100 cells per FOV. Hill fit, IC_50_ : half maximal inhibitory concentration. Two-tailed Mann-Whitney tests (A, C) and Spearman correlation (B).

Arachidonic acid (AA) is a major product of cPLA2 activity and a regulator of myosin activity (34). Deformation of the nucleus in other cell types was previously found to promote cell migration through this mechanism (16, 17). Thus, we though to assess the implication of cPLA2 in epithelial cell migration. To do so, we tracked nuclei-stained cells at confluence under increasing amounts of the cPLA2 inhibitor AACOCF3. Consistently, we found that cell migration speed decreased with AACOCF3 concentration (Fig. 5D). To further identify how cPLA2 controlled epithelial cell migration, we examined the cell shape factor, an indicator of the balance between cortical contractility and intercellular adhesion thought to control epithelial tissue jamming (35). While we found that the shape factor decreased with cell density, thereby approaching a rigidity transition (around 3.81) as cells became more packed, inhibition of cPLA2 had in contrast no visible effect (Fig.S5A). Therefore, cPLA2 does not control epithelial cell migration through cortical contractility or intercellular adhesion. To further examine how cPLA2 controlled cell migration, we measured the rate of neighbor dispersion, which accounts for the balance between migration persistence and velocity coordination between neighbors. We found that pairs of neighbors moved away form each other at about the same rate regardless of cPLA2 inhibitor concentration (Fig. S5B), indicating that cPLA2 controls cell migration speed independently of migration persistence or coordination between neighbors. Altogether, these results support that cPLA2 activity targets the core machinery of cell migration that regulates cell speed.

## Discussion

In this study, we thought to assess how epithelial cell density may control elementary cell behaviors implicated in tissue homeostasis. In summary, we found that epithelial cells respond to density through a pathway that implicates activation of ERK by mechanosensitive FAs and its translocation in the nucleus, where it feeds back to the contractile migration machinery through the activation of cPLA2 (Fig. S5C).

ERK is an central regulator of cell behavior that can be activated by a variety of extracellular signals relayed by cell surface receptors for soluble or immobilized ligands (24). Moreover, ERK activity often displays spatiotemporal fluctuations at multiple scales across tissues, in the form of stochastic pulses and waves to which each input signal may contribute in varied proportions (36). Spontaneous differences in ERK activity among the cell population are clearly visible throughout our experiments. These differences may reflect the action of all the other inputs ERK is sensitive too. Yet, the capacity of ERK to sense epithelial density appears to rely essentially on vinculin and its tension (Fig. 1A-C, S1C).

We show that low density favors FA growth, which relaxes vinculin tension (Fig. 1D, S1C,F). This might appear counterintuitive at first, since epithelial cells exert greater traction forces at the edge of a colony where density is lower, compared to packed cells in the bulk of the tissue (37). However, the growth of FAs may redistribute this traction over a larger pool of proteins, as previously observed on talin in *Drosophila* muscle cells (38), resulting in a lower tension per protein. In epithelia, mechanical changes stemming from the cell-matrix interface also impact the mechanics of Adherens Junctions (AJs), whether through force transmission or mechanotransduction (39, 40). Vinculin notably localizes at AJs, and its tension decreases in cells at lower density close to an epithelium wound (41). Since AJs mechanically regulate ERK activity (7), it is tempting to speculate that vinculin tension also controls ERK from AJs as an indirect consequence of FA growth. However, ERK activation by AJs appears to rely on an increase, rather than an decrease in their tension (42). Therefore, the density-dependence of ERK we observe is unlikely to rely on AJs. Consistently, the persistence of ERK activation by FAs growth in cells unable to transmit forces through their cytoskeleton supports a local mechanism (Fig.1D).

Interactions of vinculin with its FA partners were previously shown to regulate ERK activity (9). Specifically, paxillin serves as as a scaffold that directly interacts with ERK and FAK, and that is competed by the binding of paxillin to the tail of vinculin. We expand these findings by showing that these interactions are density-dependent (Fig. 2, S2). We propose that the growth of FAs permits a local accumulation of the proteins, and the relaxation of vinculin a conformation change that weakens its interaction with paxillin, thereby favoring the formation of a close ERK-paxillin-FAK complex (Fig. 2E). Furthermore, the nanometric vicinities we detect by FRET reveal more than known interactions and their context-dependence. At high density, the proximity of FAK and ERK with vinculin, at the expense of their mutual proximity, may reflect the competition for space in packed FAs, which persists in complexes that dynamically exchange in the cytoplasm (43). At low density, the proximity of ERK and FAK in the cytoplasm may conversely reflect the activation of FAK by ERK, that promotes its release from FAs (44). Consistently, ERK activity and its dependence on density require FAK activity (Fig. 2D). Further studies should address the dynamics of these interactions.

FA growth activates ERK, which is followed by its translocation in the nucleus within 10 minutes (Fig. 3A-B, S3A-B). Interestingly, this mode of activation appears to also result in the translocation of FAK, unlike the activation of ERK by PKC (with PMA) (Fig. S3C). In addition to its functions in FAs, FAK is also known to regulate nuclear functions (45). This opens the possibility that the combination of ERK and FAK in the nucleus results in a response specific to their activation by FAs. Additionally, FAs may regulate this response indirectly, by modulating the nucleocytoplasmic transport of ERK through nuclear deformation. Indeed, our results support a model in which the growth of FAs results in an increase of tension on LINC complexes, the disruption of which or of the cytoskeleton impairs ERK translocation (Fig.3D-G). Interestingly, the deformation of the nucleus at low density is only partially dependent on vinculin (Fig. 3E), suggesting that the other essential force transmitter of FAs talin accounts for the remaining deformation. The response is also non-linear, in agreement with the onset of NE wrinkles smoothening that occurs in this density range, and that tunes nucleocytoplasmic transport (46). The ERK-KTR sensor, which relies on nucleoplasmic transport, may owe its similarly abrupt response in part to this effect. Another possibility is that of a positive feedback loop between the nuclear translocation of ERK upon nucleus deformation and ERK activity (see below for potential mechanisms). Consistently, ERK activity at low density does not appear as high in KO cells with less deformed nuclei as in WT cells (Fig. 1A, 3E).

The lack of sensitivity of ERK to the remaining nuclear deformation in KO cells and the incomplete impairment of ERK translocation upon disruption of force transmission through LINC complexes further support that only a part of ERK nuclear translocation and activity depends on nuclear deformation, and only a part of nuclear deformation contributes to this dependence (Fig.3G, S3I). This might be related to the multiple mechanisms of ERK nuclear translocation, which leverage active transport, but also the release of cytoplasmic retention and passive transport (24). We see that the nuclear translocation of importin-7 correlates with nucleus deformation and is dependent on ERK activation (Fig. S3C,D). This is consistent with active transport acquiring mechanosensitivity through cargo-binding (47). Nevertheless, passive transport can be even more mechanosensitive than active transport (48). Future studies should therefore clarify the passive and active contributions to the mechanosensitive and insensitive parts of ERK nuclear translocation.

While the control of transcription is considered a major activity of ERK in the nucleus, we show here that ERK also functions as regulator of nuclear mechanical properties, with substantial consequences on cell behavior (Fig. 4,5). Our results support a model by which ERK-induced chromatin decondensation provides another source of nuclear envelope tension in addition to cytoskeletal forces transmitted to LINC complexes (Fig. 4). This would offer a mechanism by which ERK could facilitate its own nuclear entry by dilating nuclear pores. It may as well contribute to the deformation of the nucleus we observe at low density (Fig. 1A,3E). We propose that the effect of ERK on chromatin decondensation involves the phosphorylation-induced relocalization of Lamins to the nuclear interior that accompanies transcriptional activity (Fig. 4B, S4A), although other mechanisms, such as histone phosphorylation (49), could be at play too.

The phospholipase cPLA2 was recently found to act as a nuclear mechanotransducer upstream a behavior of cell evasion by amoeboid motility under acute nucleus confinement (16, 17). Deregulated cPLA2 activity is also known to promote epithelium-to-mesenchyme transition, migration and invasion in a variety of carcinoma cell lines (18, 19). Our results support that the control of epithelial cell migration by this nuclear mechanotransduction pathway is a physiological response to cell density (Fig. 5). We speculate that the contributions of ERK through cPLA2 phosphorylation and NE tension mediated by chromatin decondensation may compensate for the milder nuclear deformations induced by cell density compared to that experienced by cells migrating in more confined environments. Interestingly, cPLA2 was found to promote ERK phosphorylation through the Phosphoinositide 3-kinase pathway in hepatocarcinoma cells (19). This would offer a potential mechanism by which ERK could positively feedback to its activity, thereby acting as a two-state switch as a function of density, as exemplified here (Fig. 1A). As other possible positive feedback mechanisms, the effect of cPLA2 on actomyosin contractility may also promote tension on the nucleus and thereby facilitate the translocation of ERK into the nucleus.

Since arachidonic acid targets actomyosin activity, it could affect epithelial cell migration in several ways. Indeed, collective cell migration speed, persistence, and coordination depend on the combination of intrinsic and intercellular adhesion-dependent mechanisms (41), all of which involve contractile structures such as ventral stress fibers and the cortex. Our results support, however, that cPLA2 activity does not substantially affect cortical contractility (Fig. S5), which would otherwise impact the resistance of cell shape to migration (50). Consistently, neither cell persistence nor coordination, which also depend on cortex-supported intercellular adhesion, are affected (Fig. S5). Instead, our results suggest that the ventral stress fibers that are involved in the intrinsic migration machinery of the cell are targeted. Finally, we cannot exclude a contribution of perinuclear actin to nuclear deformations that would facilitate migration (51). In any case, this specificity suggests an evolutionary link to ancestral unicellular escape mechanisms under mechanical constraint, yet it questions how some structures would remain more sensitive that others in a multicellular context.

## Materials and Methods

Live MDCK type II cells transiently or stably expressing fluorescently tagged proteins or biosensors were monitored on scanning or spinning confocal epifluorescence inverted microscopes equipped for FRET, FLIM FRAP imaging and laser ablation or optical tweezer manipulation. Image analyses were performed with Fiji, CellPose, StarDist, and data analyzed with Matlab or Prism. Full materials and methods are available in Supplementary Materials and Methods. New materials, reagents, and analysis scripts are available upon request.

## Data availability

This study includes no data deposited in external repositories.

## ACKNOWLEDGEMENTS

We thank Célia Galleri–Paris, Linda Diedhiou and Thanusthika Thayatharan for preliminary experiments. We thank the members of the laboratory, Franck Riquet and Matthieu Piel for insightful discussions. We thank W. James Nelson, Martijn Gloerich, Jared Toettcher, Carsten Grashoff, Franck Riquet, Marc Tramier, Michael Davidson, Christopher E. Turner, Anna Huttenlocher and Eric Schirmer for the gift of plasmids and cell lines. This work was supported in part by the Centre national de la recherche scientifique (CNRS), the French National Research Agency (ANR) grants (ANR-17-CE13-0013, ANR-18-CE13-0008), the Investments for the Future program of the French Government (LabEx Who am I? ANR-11-LABX-0071, Université Paris Cité ANR-18-IDEX-0001) and the Fondation Recherche Médicale (EQU202203014613). We acknowledge the ImagoSeine facility at the Institut Jacques Monod, NeurImag facility at the Institute of Psychiatry and Neuroscience of Paris, members of the France BioImaging infrastructure (ANR-10-INSB-04).

## References

1. Sally P. Leys and Ana Riesgo. Epithelia, an Evolutionary Novelty of Metazoans. Journal of Experimental Zoology Part B: Molecular and Developmental Evolution, 318(6):438–447, 2012. ISSN 1552-5015. doi: 10.1002/jez.b.21442. _eprint: https://onlinelibrary.wiley.com/doi/pdf/10.1002/jez.b.21442.

2. Ian G. Macara, Richard Guyer, Graham Richardson, Yongliang Huo, and Syed M. Ahmed. Epithelial Homeostasis. Current Biology, 24(17):R815–R825, September 2014. ISSN 0960-9822. doi: 10.1016/j.cub.2014.06.068. Publisher: Elsevier.

3. Andrea I McClatchey and Alpha S Yap. Contact inhibition (of proliferation) redux. Current Opinion in Cell Biology, 24(5):685–694, October 2012. ISSN 0955-0674. doi: 10.1016/j.ceb.2012.06.009.

4. Elizabeth Lawson-Keister and M. Lisa Manning. Jamming and arrest of cell motion in biological tissues. Current Opinion in Cell Biology, 72:146–155, October 2021. ISSN 0955-0674. doi: 10.1016/J.CEB.2021.07.011. 2102.11255 Publisher: Elsevier Current Trends.

5. Lisa Donker, Ronja Houtekamer, Marjolein Vliem, François Sipieter, Helena Canever, Manuel Gómez-González, Miquel Bosch-Padrós, Willem-Jan Pannekoek, Xavier Trepat, Nicolas Borghi, and Martijn Gloerich. A mechanical G2 checkpoint controls epithelial cell division through E-cadherin-mediated regulation of Wee1-Cdk1. Cell Reports, 41(2), October 2022. ISSN 2211-1247. doi: 10.1016/j.celrep.2022.111475. Publisher: Elsevier.

6. Kazuhiro Aoki, Yohei Kondo, Honda Naoki, Toru Hiratsuka, Reina E Itoh, and Michiyuki Matsuda. Propagating Wave of ERK Activation Orients Collective Cell Migration. Developmental cell, 43(3):305–317.e5, November 2017. ISSN 1878-1551. doi: 10.1016/j.devcel.2017.10.016.

7. Tsuyoshi Hirashima, Naoya Hino, Kazuhiro Aoki, and Michiyuki Matsuda. Stretching the limits of extracellular signal-related kinase (ERK) signaling — Cell mechanosensing to ERK activation. Current Opinion in Cell Biology, 84:102217, October 2023. ISSN 0955-0674. doi: 10.1016/J.CEB.2023.102217. Publisher: Elsevier Current Trends.

8. Martin A. Schwartz, Michael D. Schaller, and Mark H. Ginsberg. Integrins: Emerging Paradigms of Signal Transduction. Annual Review of Cell and Developmental Biology, 11(Volume 11, 1995):549–599, November 1995. ISSN 1081-0706, 1530-8995. doi: 10.1146/annurev.cb.11.110195.003001. Publisher: Annual Reviews.

9. M Cecilia Subauste, Olivier Pertz, Eileen D Adamson, Christopher E Turner, Sachiko Junger, and Klaus M Hahn. Vinculin modulation of paxillin-FAK interactions regulates ERK to control survival and motility. The Journal of cell biology, 165(3):371–81, May 2004. ISSN 0021-9525. doi: 10.1083/jcb.200308011.

10. Carsten Grashoff, B.D. Hoffman, M.D. Brenner, Ruobo Zhou, Maddy Parsons, M.T. Yang, M.A. McLean, S.G. Sligar, C.S. Chen, Taekjip Ha, and others. Measuring mechanical tension across vinculin reveals regulation of focal adhesion dynamics. Nature, 466(7303): 263–266, 2010. doi: 10.1038/nature09198. Publisher: Nature Publishing Group.

11. Naoya Hino, Leone Rossetti, Ariadna Marín-Llauradó, Kazuhiro Aoki, Xavier Trepat, Michiyuki Matsuda, and Tsuyoshi Hirashima. ERK-Mediated Mechanochemical Waves Direct Collective Cell Polarization. Developmental Cell, 53(6):646–660.e8, June 2020.

12. Richard L. Klemke, Shuang Cai, Ana L. Giannini, Patricia J. Gallagher, Primal de Lanerolle, and David A. Cheresh. Regulation of Cell Motility by Mitogen-activated Protein Kinase. Journal of Cell Biology, 137(2):481–492, April 1997. ISSN 0021-9525. doi: 10.1083/jcb.137.2.481.

13. Rey-Huei Chen, Charlyn Sarnecki, and John Blenis. Nuclear Localization and Regulation of erk-and rsk-Encoded Protein Kinases. Molecular and Cellular Biology, 12(3):915–927, March 1992. ISSN null. doi: 10.1128/mcb.12.3.915-927.1992. Publisher: Taylor & Francis _eprint: https://doi.org/10.1128/mcb.12.3.915-927.1992.

14. Théophile Déjardin, Pietro Salvatore Carollo, François Sipieter, Patricia M. Davidson, Cynthia Seiler, Damien Cuvelier, Bruno Cadot, Cecile Sykes, Edgar R. Gomes, and Nicolas Borghi. Nesprins are mechanotransducers that discriminate epithelial–mesenchymal transition programs. Journal of Cell Biology, 219(10):e201908036, October 2020. ISSN 0021-9525. doi: 10.1083/jcb.201908036. Publisher: The Rockefeller University Press.

15. Alberto Elosegui-Artola, Ion Andreu, Amy E.M. Beedle, Ainhoa Lezamiz, Marina Uroz, Anita J. Kosmalska, Roger Oria, Jenny Z. Kechagia, Palma Rico-Lastres, Anabel-Lise Le Roux, Catherine M. Shanahan, Xavier Trepat, Daniel Navajas, Sergi Garcia-Manyes, and Pere Roca-Cusachs. Force Triggers YAP Nuclear Entry by Regulating Transport across Nuclear Pores. Cell, 0(0), October 2017. ISSN 00928674. doi: 10.1016/j.cell.2017.10.008. Publisher: Elsevier.

16. Valeria Venturini, Fabio Pezzano, Frederic Català Castro Hanna-Maria Häkkinen, Senda Jiménez-delgado, Mariona Colomer-rosell, Monica Marro, Queralt Tolosa-ramon, Sonia Paz-lópez, Miguel A Valverde, Julian Weghuber, Pablo Loza-alvarez, Michael Krieg, Stefan Wieser, and Verena Ruprecht. The nucleus measures shape changes for cellular proprioception to control dynamic cell behavior. Science, 370(October), October 2020. doi: 10.1126/science.aba2644. Publisher: American Association for the Advancement of Science.

17. A. J. Lomakin, C. J. Cattin, D. Cuvelier, Z. Alraies, M. Molina, GP F Nader, N. Srivastava, P. J. Saez, J. M. Garcia-Arcos, I. Y. Zhitnyak, A. Bhargava, M. K. Driscoll, E. S. Welf, R. Fi-olka, R. J. Petrie, N. S. de Silva, J.M. González-Granado, N. Manel, A.M. Lennon-Duménil, D. J. Müller, M. Piel, G. P.F. Nader, N. Srivastava, P. J. Saez, J. M. Garcia-Arcos, I. Y. Zhitnyak, A. Bhargava, M. K. Driscoll, E. S. Welf, R. Fiolka, R. J. Petrie, N. S. de Silva, J.M. González-Granado, N. Manel, A.M. Lennon-Duménil, D. J. Müller, and M. Piel. The nucleus acts as a ruler tailoring cell responses to spatial constraints. Science, 370(6514), October 2020. ISSN 10959203. doi: 10.1126/science.aba2894. Publisher: American Association for the Advancement of Science.

18. Lu Chen, Hui Fu, Yi Luo, Liwei Chen, Runfen Cheng, Ning Zhang, and Hua Guo. cPLA2α mediates TGF-β-induced epithelial–mesenchymal transition in breast cancer through PI3k/Akt signaling. Cell Death & Disease 2017 8:4, 8(4):e2728–e2728, April 2017. ISSN 2041-4889. doi: 10.1038/cddis.2017.152. Publisher: Nature Publishing Group.

19. Hui Fu, Yuchao He, Lisha Qi, Lu Chen, Yi Luo, Liwei Chen, Yongmei Li, Ning Zhang, and Hua Guo. cPLA2α activates PI3K/AKT and inhibits Smad2/3 during epithelial–mesenchymal transition of hepatocellular carcinoma cells. Cancer Letters, 403:260–270, September 2017. ISSN 0304-3835. doi: 10.1016/J.CANLET.2017.06.022. Publisher: Elsevier.

20. François Sipieter, Benjamin Cappe, Mariano Gonzalez Pisfil, Corentin Spriet, Jean-François Bodart, Katia Cailliau-Maggio, Peter Vandenabeele, Laurent Héliot, and Franck B. Riquet. Novel Reporter for Faithful Monitoring of ERK2 Dynamics in Living Cells and Model Organisms. PLOS ONE, 10(10):e0140924, October 2015. ISSN 1932-6203. doi: 10.1371/journal.pone.0140924. Publisher: Public Library of Science.

21. James A. Lorenzen, Scott E. Baker, Fabienne Denhez, Michael B. Melnick, Danny L. Brower, and Lizabeth A. Perkins. Nuclear import of activated D-ERK by DIM-7, an importin family member encoded by the gene moleskin. Development, 128(8):1403–1414, April 2001. ISSN 0950-1991. doi: 10.1242/dev.128.8.1403.

22. Kylie M. Wagstaff, Haran Sivakumaran, Steven M. Heaton, David Harrich, and David A. Jans. Ivermectin is a specific inhibitor of importin α/β-mediated nuclear import able to inhibit replication of HIV-1 and dengue virus. Biochemical Journal, 443(3):851–856, April 2012. ISSN 0264-6021. doi: 10.1042/BJ20120150.

23. GW Gant Luxton, Edgar R Gomes, Eric S Folker, Erin Vintinner, and Gregg G Gundersen. Linear arrays of nuclear envelope proteins harness retrograde actin flow for nuclear movement. Science (New York, N.Y.), 329(5994):956–9, August 2010. ISSN 1095-9203. doi: 10.1126/science.1189072.

24. Hugo Lavoie, Jessica Gagnon, and Marc Therrien. ERK signalling: a master regulator of cell behaviour, life and fate. Nature Reviews Molecular Cell Biology 2020 21:10, 21(10): 607–632, June 2020. ISSN 1471-0080. doi: 10.1038/s41580-020-0255-7. Publisher: Nature Publishing Group.

25. Nicolas Audugé, Sergi Padilla-Parra, Marc Tramier, Nicolas Borghi, and Maïté Coppey-Moisan. Chromatin condensation fluctuations rather than steady-state predict chromatin accessibility. Nucleic Acids Research, 47(12):6184–6194, May 2019. ISSN 0305-1048. doi: 10.1093/nar/gkz373. Publisher: Oxford Academic.

26. Kohta Ikegami, Stefano Secchia, Omar Almakki, Jason D. Lieb, and Ivan P. Moskowitz. Phosphorylated Lamin A/C in the Nuclear Interior Binds Active Enhancers Associated with Abnormal Transcription in Progeria. Developmental Cell, 52(6):699–713.e11, March 2020. ISSN 1534-5807. doi: 10.1016/j.devcel.2020.02.011. Publisher: Elsevier.

27. Scott M. Carlson, Candace R. Chouinard, Adam Labadorf, Carol J. Lam, Katrin Schmelzle, Ernest Fraenkel, and Forest M. White. Large-Scale Discovery of ERK2 Substrates Identifies ERK-Mediated Transcriptional Regulation by ETV3. Science Signaling, 4(196):rs11–rs11, October 2011. doi: 10.1126/scisignal.2002010. Publisher: American Association for the Advancement of Science.

28. Vitaly Kochin, Takeshi Shimi, Elin Torvaldson, Stephen A. Adam, Anne Goldman, Chan-Gi Pack, Johanna Melo-Cardenas, Susumu Y. Imanishi, Robert D. Goldman, and John E. Eriksson. Interphase phosphorylation of lamin A. Journal of Cell Science, 127(12):2683– 2696, June 2014. ISSN 0021-9533. doi: 10.1242/jcs.141820.

29. Aprotim Mazumder, T. Roopa, Aakash Basu, L. Mahadevan, and G. V. Shivashankar. Dynamics of Chromatin Decondensation Reveals the Structural Integrity of a Mechanically Prestressed Nucleus. Biophysical Journal, 95(6):3028–3035, September 2008. ISSN 0006-3495, 1542-0086. doi: 10.1529/biophysj.108.132274. Publisher: Elsevier.

30. F. Brochard-Wyart, P.G. de Gennes, and O. Sandre. Transient pores in stretched vesicles: role of leak-out. Physica A: Statistical Mechanics and its Applications, 278(1):32–51, 2000. ISSN 03784371. doi: 10.1016/S0378-4371(99)00559-2.

31. Charlotte Alibert, David Pereira, Nathan Lardier, Sandrine Etienne-Manneville, Bruno Goud, Atef Asnacios, and Jean-Baptiste Manneville. Multiscale rheology of glioma cells. Biomaterials, 275:120903, August 2021. ISSN 0142-9612. doi: 10.1016/j.biomaterials.2021.120903.

32. Balázs Enyedi, Mark Jelcic, and Philipp Niethammer. The Cell Nucleus Serves as a Mechanotransducer of Tissue Damage-Induced Inflammation. Cell, 165(5):1160–1170, May 2016. ISSN 0092-8674. doi: 10.1016/J.CELL.2016.04.016. Publisher: Cell Press.

33. Lih Ling Lin, Markus Wartmann, Alice Y. Lin, John L. Knopf, Alpna Seth, and Roger J. Davis. cPLA2 is phosphorylated and activated by MAP kinase. Cell, 72(2):269–278, January 1993. ISSN 0092-8674. doi: 10.1016/0092-8674(93)90666-E. Publisher: Cell.

34. Ming Cui Gong, Annette Fuglsang, Dario Alessi, Sei Kobayashi, Philip Cohen, Avril V. Somlyo, and Andrew P. Somlyo. Arachidonic acid inhibits myosin light chain phosphatase and sensitizes smooth muscle to calcium. Journal of Biological Chemistry, 267(30):21492– 21498, October 1992. ISSN 0021-9258. doi: 10.1016/S0021-9258(19)36636-0. Publisher: Elsevier.

35. Dapeng Bi, J. H. Lopez, J. M. Schwarz, and M. Lisa Manning. A density-independent rigidity transition in biological tissues. Nature Physics, 11(12):1074–1079, December 2015. ISSN 1745-2473. doi: 10.1038/nphys3471. Publisher: Nature Publishing Group.

36. Kazuhiro Aoki, Yuka Kumagai, Atsuro Sakurai, Naoki Komatsu, Yoshihisa Fujita, Clara Shionyu, and Michiyuki Matsuda. Stochastic ERK activation induced by noise and cell-to-cell propagation regulates cell density-dependent proliferation. Molecular cell, 52(4):529–40, November 2013. ISSN 1097-4164. doi: 10.1016/j.molcel.2013.09.015.

37. Olivia du Roure, Alexandre Saez, Axel Buguin, Robert H Austin, Philippe Chavrier, Pascal Silberzan, and Benoit Ladoux. Force mapping in epithelial cell migration. Proceedings of the National Academy of Sciences, 102(7):2390–2395, February 2005.

38. Sandra B. Lemke, Thomas Weidemann, Anna-Lena Cost, Carsten Grashoff, and Frank Schnorrer. A small proportion of Talin molecules transmit forces at developing muscle attachments in vivo. PLOS Biology, 17(3):e3000057, March 2019. ISSN 1545-7885. doi: 10.1371/journal.pbio.3000057. Publisher: Public Library of Science.

39. Xavier Trepat, Michael R Wasserman, Thomas E Angelini, Emil Millet, David A Weitz, James P Butler, and Jeffrey J Fredberg. Physical forces during collective cell migration. Nature Physics, 5(6):426–430, 2009. ISSN 1745-2473. doi: 10.1038/nphys1269. Publisher: Nature Publishing Group.

40. Charlène Gayrard, Clément Bernaudin, Théophile Déjardin, Cynthia Seiler, and Nicolas Borghi. Src- and confinement-dependent FAK activation causes E-cadherin relaxation and β-catenin activity. The Journal of cell biology, 217(3):1063–1077, March 2018. ISSN 1540-8140. doi: 10.1083/jcb.201706013. Publisher: Rockefeller University Press.

41. Helena Canever, Hugo Lachuer, Quentin Delaunay, François Sipieter, Nicolas Audugé, Philippe P. Girard, and Nicolas Borghi. Collective directional memory controls the range of epithelial cell migration, January 2025. Pages: 2023.12.08.570811 Section: New Results.

42. Ronja M. Houtekamer, Mirjam C. van der Net, Marjolein J. Vliem, Tomas E. J. C. Noordzij, Lisa van Uden, Robert M. van Es, Joo Yong Sim, Eriko Deguchi, Kenta Terai, Matthew A. Hopcroft, Harmjan R. Vos, Beth L. Pruitt, Michiyuki Matsuda, Willem-Jan Pannekoek, and Martijn Gloerich. E-cadherin mechanotransduction activates EGFR-ERK signaling in epithelial monolayers by inducing ADAM-mediated ligand shedding. Science Signaling, 18 (886):eadr7926, May 2025. doi: 10.1126/scisignal.adr7926. Publisher: American Association for the Advancement of Science.

43. J.-E. Hoffmann, Y. Fermin, R. L. Stricker, K. Ickstadt, and E. Zamir. Symmetric exchange of multi-protein building blocks between stationary focal adhesions and the cytosol. eLife, 3:e02257–e02257, June 2014. ISSN 2050-084X. doi: 10.7554/eLife.02257. Publisher: eLife Sciences Publications Limited.

44. Satyajit K Mitra, Daniel A Hanson, and David D Schlaepfer. Focal adhesion kinase: in command and control of cell motility. Nature reviews. Molecular cell biology, 6(1):56–68, January 2005. ISSN 1471-0072. doi: 10.1038/nrm1549.

45. Elizabeth G Kleinschmidt and David D Schlaepfer. Focal adhesion kinase signaling in unexpected places. Current Opinion in Cell Biology, 45:24–30, April 2017. ISSN 0955-0674. doi: 10.1016/j.ceb.2017.01.003.

46. Ignasi Granero-Moya, Valeria Venturini, Guillaume Belthier, Bart Groenen, Marc Molina-Jordán, Miguel González-Martín, Xavier Trepat, Jacco van Rheenen, Ion Andreu, and Pere Roca-Cusachs. Nucleocytoplasmic transport senses mechanical forces independently of cell density in cell monolayers. Journal of Cell Science, 137(17):jcs262363, September 2024. ISSN 0021-9533. doi: 10.1242/jcs.262363.

47. María García-García, Sara Sánchez-Perales, Patricia Jarabo, Enrique Calvo, Trevor Huyton, Liran Fu, Sheung Chun Ng, Laura Sotodosos-Alonso, Jesús Vázquez, Sergio Casas-Tintó, Dirk Görlich, Asier Echarri, and Miguel A. Del Pozo. Mechanical control of nuclear import by Importin-7 is regulated by its dominant cargo YAP. Nature Communications 2022 13:1, 13(1):1–21, March 2022. ISSN 2041-1723. doi: 10.1038/s41467-022-28693-y. Publisher: Nature Publishing Group.

48. Ion Andreu, Ignasi Granero-Moya, Nimesh R. Chahare, Kessem Clein, Marc Molina-Jordán, Amy E. M. Beedle, Alberto Elosegui-Artola, Juan F. Abenza, Leone Rossetti, Xavier Trepat, Barak Raveh, Pere Roca-Cusachs, Harvard A John Paulson, Ion Andreu, Ignasi Granero-Moya, Nimesh R. Chahare, Kessem Clein, Marc Molina-Jordán, Amy E M Beedle, Alberto Elosegui-Artola, Juan F. Abenza, Leone Rossetti, Xavier Trepat, Barak Raveh, and Pere Roca-Cusachs. Mechanical force application to the nucleus regulates nucleocytoplasmic transport. Nature Cell Biology, (24):896–905, June 2022. ISSN 1476-4679. doi: 10.1038/s41556-022-00927-7. Publisher: Nature Publishing Group.

49. Deborah N. Chadee, Michael J. Hendzel, Cheryl P. Tylipski, C. David Allis, David P. Bazett-Jones, Jim A. Wright, and James R. Davie. Increased Ser-10 Phosphorylation of Histone H3 in Mitogen-stimulated and Oncogene-transformed Mouse Fibroblasts *. Journal of Biological Chemistry, 274(35):24914–24920, August 1999. ISSN 0021-9258, 1083-351X. doi: 10.1074/jbc.274.35.24914. Publisher: Elsevier.

50. Dapeng Bi, Xingbo Yang, M. Cristina Marchetti, and M. Lisa Manning. Motility-driven glass and jamming transitions in biological tissues. Physical Review X, 6(2):021011, April 2016. ISSN 21603308. doi: 10.1103/PhysRevX.6.021011. 1509.06578 Publisher: American Physical Society.

51. Patricia M. Davidson and Bruno Cadot. Actin on and around the Nucleus. Trends in Cell Biology, December 2020. ISSN 09628924. doi: 10.1016/j.tcb.2020.11.009. Publisher: Elsevier Current Trends.

