## Supplementary material for "Epithelial density controls cell migration through an adhesion-nucleus mechanotransduction pathway": SI M&M, tables and figures

### Materials and Methods

**Cell lines, plasmids and sample preparation.** The parental Madin Darby Canine Kidney (MDCK) type II cell line was a gift from W. James Nelson. All generated cell lines were cultured at 37°C and 5% CO2 in Dulbecco's Modified Eagle's Medium (DMEM) with low D-glucose content (1g/l) containing Pyruvate and L-Glutamine (Gibco, 31885-023), supplemented with 10% Fetal Bovine Serum (FBS) (Gibco, 10270-106), 10 U / ml Penicillin, 10 µg/ml Streptomycin (Gibco, 15140-122), and 1% (vol / vol) L-glutamine (Gibco, 250). Cell lines expressing the mini-Nesprin-2 Giant (mN2G) CB and CH constructs (1) were maintained in DMEM with 100$\mu$g/mL Hygromycin. The Vinculin$^{-/-}$ cell line is described elsewhere (2).
Cells were transfected with TurboFect (R0531, Thermo Fisher Scientific) according to the manufacturer's recommendations with the following plasmids:

- ERK-KTR. To construct ERK-KTR-iRFP713 (NeoR and HygroR), we used PCR and restriction cloning (NheI/BsrGI and NheI/AgeI, respectively) to insert ERK-KTR-iRFP713 (from Addgene 111510, a gift from Jared Toettcher) into pmEGFP-N1 (Clontech, NeoR) and pcDNA3.1(+) (Thermo, HygroR), respectively.
- VinTS (a gift from Carsten Grashoff, (3)). The V1001A mutant was generated from this construct by site-directed mutagenesis (QuikChange II XL Kit, Agilent Technologies) with primers 5'-CCCAGCATGGTAGCTTTCGCTGTGGAAAGAATTTTGA-3' and 5'-TCAAAATTCTTTCCACAGCGAAAGCTACCATGCTGGG-3', as well as the T12 mutant described elsewhere (4).
- EKAR. To create EKAR-GW4.0_NLS (YPET/mTFP1), eCFP in pCS2-EKAR-GW4.0 (YPET/eCFP) (a gift from Franck Riquet (5)) was replaced with mTFP1-NLS by PCR amplification using pmTFP1-ShadowG as template (a gift from Marc Tramier, Addgene 99867), PCR primers fused to PspXI and NotI-NLS, and next subcloning using PspXI and NotI restriction. The resulting EKAR-GW4.0_NLS (YPET/mTFP1) was subsequently subcloned into pmEGFP-N1 (NeoR) via BamHI and NotI restriction sites. For multiplexed biosensing experiments, we replaced YPET and eCFP in pCS2-EKAR-GW4.0 (YPET/eCFP) with mKate2 (from pLSSmOrange-mKate2, a gift from Marc Tramier, Addgene 99868) and mRuby2 (from Clover-mRuby2 tandem, a gift from Michael Davidson, Addgene 58169), respectively, by PCR amplification, restriction digest and ligation, resulting in pCS2-EKAR-GW4.0 (mKate2/mRuby2). For mKate2 and mRuby2 PCR amplifications, we used PCR primers fused to BamHI/BsrZ17I and PspXI/NotI restriction sites, respectively. We next subcloned the EKAR-GW4.0 (mKate2/mRuby2) into pcDNA3.1(+) (HygroR) via BamHI and NotI restriction sites.
- VCL-eGFP. To create this construct, we replaced mCherry in mCherry-Vinculin-N-21 (a gift from Michael Davidson, Addgene 55160) with mEGFP (pmEGFP-N1, Clontech) by restriction digest (EcoRI/NotI) and ligation, resulting in meGFP-Vinculin-N-21 (NeoR).
- eGFP-PXL (pmEGFP-C1-paxillin, a gift from Christopher E. Turner (6)).
- mCherry-PXL. We replaced mEGFP in pmEGFP-C1-paxillin with mCherry (pmCherry-C1, Clontech) by restriction digest (NheI/BspEI) and ligation, resulting in pmCherry-C1-paxillin.
- FAK-eGFP, FAK-mCherry. FAK was PCR amplified using pmCherry-C1-FAK-HA as a template (a gift from Anna Huttenlocher, Addgene 35039) and PCR primers fused to NheI and SalI restriction sites. FAK was then subcloned into pmCherry-N1 and pmEGFP-N1 (Clontech) by restriction digest (NheI/SalI) and ligation, resulting in pmCherry-N1-FAK and pmEGFP-N1-FAK, respectively.
- ERK2Loc (a gift from Franck Riquet (7)).
- mCherry-ERK2. We replaced eGFP in ERK2Loc with mCherry (pmCherry-C1, Clontech) by restriction digest (AgeI/BsrGI) and ligation.
- DNKASH-mCherry (1).
- LMNA-mRFP (a gift from Eric Schirmer, Addgene 124268).

DNA fragments of new constructions were all purified on Qiagen plasmid purification columns (28106, 28706 and 27106, Qiagen). All resulting constructs were verified by restriction digestion followed by agarose gel electrophoresis, or by PCR colony screening (#2200210, MasterTaq Kit, 5Prime), and then validated by sequencing (Eurofins Genomics). Stable cell lines expressing ERK-KTR, VinTS, V1001A, T12, EKAR, ERK2Loc, mCherry-ERK2 or LMNA-mRFP were obtained by FACS and maintained with 200 µg/ml Geneticin (G418) (Gibco, 10131-035) or 100 µg/ml Hygromycin B Gold (Invivogen, ant-hg).
For all experiments, cells were washed with Dulbecco's Phosphate Buffered Saline (DPBS) 1X (Gibco, 14190-094) and detached with a 0.05% Trypsin solution with Ethylene Diamine Tetra-Acetic Acid (EDTA) and Phenol Red (Gibco, 25300-054) for 15 minutes, counted with a Countess 3 Automated Cell Counter (ThermoFisher Scientific). Suspensions of $1.2.10^5$

cells/mL for high density, and $6.10^3$ cells/ml for low density, were deposited 48 hours before acquisition in DMEM FluoroBriteTM (Gibco, A18967-01) with 2.5 mM L-Glutamine and supplemented with 1 or 10% (vol/vol) FBS in compartmentalized glass-bottom slides with 2, 4, or 8 wells NuncTM Lab-TekTM (ThermoFisher Scientific, 155383) or µ-Slide ibiTreat (Ibidi, 80287/80421/80806) previously coated by immersion for 1 hour in a solution of 50 µg/ml human placental Collagen IV (Sigma-Aldrich, C5533) solubilized in 0.5 M glacial acetic acid (Avantor VWR, 20104.298) followed by exposure to UV rays for 15 minutes and two 5-minute washes with DPBS 1X.

**Pharmacological perturbations.** Cells were treated as follows: 20 minutes at 1 mM for $Mn^{2+}$ (Manganese chloride, Sigma-Aldrich, M1787); 20 minutes at 1 µM for GLP (GLPG0187, MedChemExpress, HY-100506); 20 minutes at 0.5 µM for PMA (Phorbol 12-Myristate 13-Acetate, Sigma-Aldrich, P8139); 20 minutes at 10 µM for SCH (SCH772984, Selleckchem, S7101) and PF228 (PF-573228, Selleckchem, S2013); 15 minutes at 0.5 µM for cytoD (cytochalasin D, MilliporeSigma, C8273); 30 minutes at 1,10 or 20 µM for AACOCF3 (Arachidonyl trifluoromethyl ketone, Abcam, AB120350); and 1 hour at 25 µM for Ivermectin (Sigma-Aldrich, I8898).

**FRET biosensors imaging and analysis.** Spectral imaging was performed on live cells at 37°C and 5% CO2 on a confocal microscope (Carl Zeiss LSM 780) with a 63×/1.4 NA Plan-Apochromat oil immersion objective. Blue/yellow sensors were excited by the 458-nm line of a 25-mW argon laser and emission was sampled at a spectral resolution of 8.7 nm within the 476–557-nm range on a GaAsP detector. Red/far-red sensors were excited by the 561-nm line of a 15-mW DPSS laser and emission was then sampled in the 571–664 nm range. For time-lapse experiments, images were acquired every 2 min during 20 min. Image analysis and FRET index measurement was performed on Fiji 1.52 or 1.53t with the open-source PixFRET plugin or with a home-made script described elsewhere for mN2G sensors (8). The FRET index was defined as $I_{\mathrm{A}}/(I_{\mathrm{D}}+I_{\mathrm{A}})$, where $I_{\mathrm{A}}$ and $I_{\mathrm{D}}$ are the background-subtracted donor and acceptor intensities, respectively, upon donor excitation.

**FLIM.** FLIM-FRET experiments on FA proteins were conducted on live cells at 37°C and 5% CO2 on a Leica SP8 Falcon system equipped with a hybrid detector for time-correlated single-photon counting (Leica Microsystems, Germany). eGFP was excited at 488 nm with a white light pulsed laser (50 MHz) and emission was collected from 496 nm to 536 nm through a 63X/1.4 NA oil objective with apochromatic correction. Fluorescence lifetimes were determined in the LAS X software (Leica Microsystems) by fitting a mono-exponential decay model to the photon count histograms ($\chi^2$ < 1.0, fitting range 0.048 ns to 12.364 ns) built from all the pixels that collected at least 50 photons each within regions of interest (Focal Adhesions or cytoplasm) defined from the corresponding intensity image.
FLIM on H4-GFP was performed on a system that combines multifocal multiphoton excitation and a fast-gated CCD camera controlled by the IMspector software (LaVisionBiotec, Bielefeld, Germany), as described elsewhere (9). Briefly, two-photon multifocal excitation was carried out with a mode-locked Ti:Sa laser at 950 nm (MaiTai, Spectra Physics, Evry, France), split into 64 beams with a 50/50 beam splitter and mirrors. The beams passed through a 2000 Hz scanner before illuminating the back aperture of an infrared water immersion objective (60×/NA 1.2, Olympus) of an inverted microscope (IX 71, Olympus). A line of foci at the focal plane was scanned across the sample, generating a pseudo wide-field illumination. Fluorescence is imaged through an emission filter (520DF40, Chroma Technology) onto a fast-gated light intensifier connected to a CCD camera (PicoStar). The intensifier applied five 2ns gates from 0 to 10 ns along the fluorescence decay. The mean, instantaneous fluorescence lifetime is defined as $\tau = \Sigma I_i \Delta t_i / \Sigma I_i$, where $\Delta t_i$ is the time delay after the laser pulse of the $i$th gate and $I_i$ the corresponding pixel intensity map. The acquisition time was set to 3s and $\tau$ was averaged over 100 consecutive images.

**Laser ablation experiments and analysis.** Laser ablation of the nuclear envelope was conducted on live cells stably expressing LMNA-mRFP at 37°C and 5% CO2 on a Yokogawa Spinning Disk CSU-X1 microscope. Cells were imaged with a 100x/1.4NA oil immersion objective. Fluorescence was acquired by 561nm laser excitation with a 590/35 emission filter and a sCMOS PRIME 95 camera (Teledyne-Photometrics). Ablation was achieved by a 355nm laser controlled via Metamorph imaging software with user-defined ROI and pulse duration settings. Time-lapse imaging was performed at 250 ms intervals over 5 minutes. Image stacks were processed with a custom script involving segmentation of the nucleus cross-section by CellPose-SAM (10) and quantification of its area $A$ in Fiji. Assuming small deflation on short time scales, $\frac{A}{A_0}(t) \simeq 1 - \frac{\pi}{3}\frac{\sigma}{\eta}\frac{r^3}{A_0^2}t$, where $A_0$ is the area before ablation, $\sigma$ the NE tension relaxed by ablation, $\eta$ the viscosity of the leaking fluid and $r$ the radius of the hole (11). To estimate $\sigma$, $A/A_0$ was therefore fit with a straight line through (0,1) for $t \leq 2.5$s, where $A_0$ and $r$ were determined from images and $\eta$ from optical tweezer experiments (see below).

**Optical tweezer experiments and analysis.** LMNA-mRFP cells were incubated with polystyrene beads of radius $r_{\mathrm{b}} = 1\mu$m (Invitrogen) 48hrs before experiments to let beads internalize. The optical tweezer setup was described previously (12). In brief, a single fixed optical trap was generated from a 1060-1100 nm infrared laser beam (2 W maximum output power; IPG Photonics) connected the back port of an inverted Eclipse microscope (Nikon) equipped with a laser confocal A1R resonant scanner (Nikon), a 37 °C incubator, and a nanometric piezostage (Mad City Labs). The nuclear indentation protocol and analysis with a custom Matlab script were described previously (13). Briefly, a bead initially contacting the nucleus was trapped with the laser and the piezostage was moved at constant speed ($2.5\mu$m/min) to push the nucleus towards the bead. The force was determined from the displacement of the bead from the trap center and the trap stiffness (240pN/$\mu$m), and the indentation depth $\delta$ was obtained by image analysis at a temporal resolution of 130ms. A Kelvin-Voigt model was used to determine the stiffness $k$ of the nucleus from the force-indentation curves (13). Then, when the displacement reached the limit of the trap, the trap released the bead and the indentation relaxed from its maximum $\delta_{\text{max}}$ towards the initial shape of the nucleus. Assuming a viscous drag on the half bead surface indented into the nucleus, the relaxation of the indentation depth was fitted with a visco-elastic exponential decay given by $\delta(t) = \delta_{\text{max}}\exp\frac{-k}{3\pi r_b \eta}t$ to determine the viscosity of the nucleus $\eta$, with $k$ obtained from the indentation phase as described above.

**Confocal imaging.** All other live and fixed fluorescence acquisitions were carried out on a Carl Zeiss LSM 780, at 37°C and 5% CO2 when necessary. Depending on their spectral properties, the fluorophores were excited with the 405 nm line using a 405-30 diode laser source; at 488 nm with an Argon laser source; at 561 nm with a 15 mW DPSS source and at 633 nm with a 5 mW Helium-Neon laser source through a 63X/1.4 NA oil objective with apochromatic correction, unless otherwise indicated. Fluorescence emission was collected on a photomultiplier detection tube over the ranges 418-507 nm for $\lambda_{\text{ex}}$= 405 nm, 493-575 nm for $\lambda_{\text{ex}}$= 488 nm, 566-614 nm for $\lambda_{\text{ex}}$ = 561 nm, and 644-758 nm for $\lambda_{\text{ex}}$= 633 nm. For time-lapse experiments after drug treatment, images were acquired every 2 min during 20 min. For FRAP experiments, photobleaching of LMNA-mRFP was performed at 100% 561nm laser power in a small ROI at the nucleus periphery and images were acquired every 20s.

**Immunofluorescence.** Cells were fixed in 4% (vol/vol) PFA in PBS (ThermoFisher Scientific, J61899.AK) for 10 minutes at room temperature, followed by a 5-minute wash in 1X PBS with 0.1M Glycine (Promega, H5073), and two 5-minute washes in 1X PBS. Cells were permeabilized with 0.1% (vol/vol) Triton X-100 (Sigma-Aldrich, T9284) in PBS for 5 minutes, followed by three 5-minute washes in 1X PBS, and incubation in 2% (wt/vol) Bovine Serum Albumin (BSA) in PBS for 30 minutes at room temperature. Cells were incubated in the BSA solution for 24 hours at room temperature with anti- pSer22-Lamin A/C (rabbit, Cell Signalling 13448), Lamin A/C (mouse, Cell Signalling 4777), pSer505-cPLA2 (rabbit, Cell Signalling 2831), cPLA2 (mouse, Santa Cruz sc-454) and Importin-7 (rabbit (Invitrogen, PA5-21764) primary antibodies, followed by three 5-minute washes with a 0.5% (vol/vol) Tween20 (Sigma-Aldrich, P1379) in PBS. Cells were incubated with the following secondary antibodies from goat in the BSA solution for 1 hour at 37°C: anti-mouse DyLight 488 (Invitrogen 35502), anti-rabbit DyLight 594 (Invitrogen 35560), followed with three 5 minutes washes with the Tween20 solution and 10 minutes incubation with 20 µM Hoechst 33342 (ThermoFisher Scientific, 62249) solution in PBS. Cell were finally mounted in Fluoromount (Sigma-Aldrich F4680).

**Fluorescence quantifications.** Nucleus segmentation was performed in Fiji with a home-made Jython script using StarDist2D (14) to segment nuclei. Corresponding cells are determined by Voronoi tessellation from nuclei centroids. Cytosolic regions are defined as a user-specified band surrounding corresponding nuclear regions within the Voronoi boundaries. Nucleo-cytoplasmic intensity ratios are defined as $I_{\text{N}}/I_{\text{C}}$ for ERK2Loc, Imp7 and pSer505-PLA2 or $I_{\text{C}}/I_{\text{N}}$ for ERK-KTR, where $I_{\text{N}}$ and $I_{\text{C}}$ are the background-subtracted intensities in the nuclear and cytosolic regions, respectively. For time lapse experiments, ratios were normalized to their initial values, and to that of the control at every time point (DMSO). Lamin turnover in FRAP experiments was assessed by normalizing its background-subtracted intensity to that of the corresponding ROI averaged over all time points before photobleaching, while correcting for global photobleaching, all with a home-made script in Fiji. cPLA2 recruitment to the nuclear envelope was measured by manually selecting nuclear envelope and nucleosol regions. The ratio is defined as $I_{\text{NE}}/I_{\text{Nu}}$ where $I_{\text{NE}}$ and $I_{\text{Nu}}$ are the background-subtracted intensities of cPLA2 labeling at the nuclear envelope and in the nucleosol, respectively.

**Migration assays.** Cells were seeded at $2.5.10^5$ cells/ml in DMEM FluoroBrite supplemented with 10% FBS on collagen-coated slides as described above 24 hours before acquisition. Cells were exposed to AACOCF3 (1, 10 or 20 µM) and 1 µM Hoechst 33342 for 30 minutes before imaging by excitation at $\lambda_{\text{ex}}$= 405 nm every 10 minutes for 2 hours through a Fluar 10X/0.5 NA air objective. Nucleus tracking was performed with TrackMate 7.10.1 (15) in Fiji 1.53t. The mean speed is defined as the average of instantaneous displacements within 10 min over whole tracks and was fit with the Hill equation $v = v_{\text{max}}/(1+[AACOFF3]/\text{IC}_{50})$. The shape index is defined as $p/\sqrt{A}$ where $p$ and $A$ are the perimeter and area of Voronoi cells generated from nucleus positions with a home-made script in Fiji as indicated above. The neighbor dispersion is defined as the average over $t$ of $D(\Delta t) = \frac{1}{n}\Sigma_{i\neq j}(|\mathbf{r}_i(\Delta t+t) - \mathbf{r}_j(\Delta t+t)| - |\mathbf{r}_i(t) - \mathbf{r}_j(t)|)$ where $\mathbf{r}_{i,j}$ are the positions of cells $i,j$ such that $|\mathbf{r}_i(t) - \mathbf{r}_j(t)| < 10\mu$m, and was obtained from nucleus tracks with a home-made Matlab script.

**Statistical analysis.** Data are represented as Mean ± Standard Error of the Mean (SEM). Statistical analyses were performed in GraphPad Prism or Python, and as indicated in figure legends. Briefly, conditions were compared with non-parametric, two-tailed Mann-Whitney or Kruskall-Wallis and Dunn's tests assuming independence between ROIs/cells/FOVs. Correlations were assessed using Pearson's, Spearman's or RM Kendall's coefficients. Slopes of linear fits were compared with the extra sum-of-squares F test with equal slope as the null hypothesis.

### References


1. Théophile Déjardin, Pietro Salvatore Carollo, François Sipieter, Patricia M. Davidson, Cynthia Seiler, Damien Cuvelier, Bruno Cadot, Cecile Sykes, Edgar R. Gomes, and Nicolas Borghi. Nesprins are mechanotransducers that discriminate epithelial–mesenchymal transition programs. *Journal of Cell Biology*, 219(10):e201908036, October 2020. ISSN 0021-9525. doi: 10.1083/jcb.201908036. Publisher: The Rockefeller University Press.
2. Jooske L. Monster, Lisa Donker, Marjolein J. Vliem, Zaw Win, Helen K. Matthews, Joleen S. Cheah, Soichiro Yamada, Johan de Rooij, Buzz Baum, and Martijn Gloerich. An asymmetric junctional mechanoresponse coordinates mitotic rounding with epithelial integrity. *Journal of Cell Biology*, 220(5), May 2021. ISSN 0021-9525. doi: 10.1083/jcb.202001042. Publisher: The Rockefeller University Press.
3. Carsten Grashoff, B.D. Hoffman, M.D. Brenner, Ruobo Zhou, Maddy Parsons, M.T. Yang, M.A. McLean, S.G. Sligar, C.S. Chen, Taekjip Ha, and others. Measuring mechanical tension across vinculin reveals regulation of focal adhesion dynamics. *Nature*, 466(7303):263–266, 2010. doi: 10.1038/nature09198. Publisher: Nature Publishing Group.
4. Helena Canever, Hugo Lachuer, Quentin Delaunay, François Sipieter, Nicolas Audugé, Philippe P. Girard, and Nicolas Borghi. Collective directional memory controls the range of epithelial cell migration, January 2025. Pages: 2023.12.08.570811 Section: New Results.
5. François Sipieter, Benjamin Cappe, Aymeric Leray, Elke De Schutter, Jolien Bridelance, Paco Hulpiau, Guy Van Camp, Wim Declercq, Laurent Héliot, Pierre Vincent, Peter Vandenabeele, and Franck B. Riquet. Characteristic ERK1/2 signaling dynamics distinguishes necroptosis from apoptosis. *iScience*, 24(9):103074, September 2021. ISSN 2589-0042. doi: 10.1016/J.ISCI.2021.103074. Publisher: Elsevier.
6. Valérie Petit, Brigitte Boyer, Delphine Lentz, Christopher E. Turner, Jean Paul Thiery, and Ana M. Vallés. Phosphorylation of Tyrosine Residues 31 and 118 on Paxillin Regulates Cell Migration through an Association with Crk in Nbt-II Cells. *Journal of Cell Biology*, 148(5):957–970, March 2000. ISSN 0021-9525. doi: 10.1083/jcb.148.5.957.
7. François Sipieter, Benjamin Cappe, Mariano Gonzalez Pisfil, Corentin Spriet, Jean-François Bodart, Katia Cailliau-Maggio, Peter Vandenabeele, Laurent Héliot, and Franck B. Riquet. Novel Reporter for Faithful Monitoring of ERK2 Dynamics in Living Cells and Model Organisms. *PLOS ONE*, 10(10):e0140924, October 2015. ISSN 1932-6203. doi: 10.1371/journal.pone.0140924. Publisher: Public Library of Science.
8. François Sipieter, Louis Laurent, Philippe P. Girard, and Nicolas Borghi. Molecular tension microscopy of the LINC complex in live cells. *STAR protocols*, 3(3):101538, September 2022. ISSN 2666-1667. doi: 10.1016/J.XPRO.2022.101538. Publisher: STAR Protoc.
9. Sergi Padilla-Parra, Nicolas Audugé, Maïté Coppey-Moisan, and Marc Tramier. Quantitative FRET analysis by fast acquisition time domain FLIM at high spatial resolution in living cells. *Biophysical journal*, 95(6):2976–2988, October 2008. ISSN 1542-0086. doi: 10.1529/biophysj.108.131276.
10. Marius Pachitariu, Michael Rariden, and Carsen Stringer. Cellpose-SAM: superhuman generalization for cellular segmentation, May 2025. Pages: 2025.04.28.651001 Section: New Results.
11. F. Brochard-Wyart, P.G. de Gennes, and O. Sandre. Transient pores in stretched vesicles: role of leak-out. *Physica A: Statistical Mechanics and its Applications*, 278(1):32–51, 2000. ISSN 03784371. doi: 10.1016/S0378-4371(99)00559-2.
12. David Guet, Kalpana Mandal, Mathieu Pinot, Jessica Hoffmann, Yara Abidine, Walter Sigaut, Sabine Bardin, Kristine Schauer, Bruno Goud, and Jean-Baptiste Manneville. Mechanical Role of Actin Dynamics in the Rheology of the Golgi Complex and in Golgi-Associated Trafficking Events. *Current Biology*, 24(15):1700–1711, August 2014. ISSN 0960-9822. doi: 10.1016/j.cub.2014.06.048. Publisher: Elsevier.
13. Charlotte Alibert, David Pereira, Nathan Lardier, Sandrine Etienne-Manneville, Bruno Goud, Atef Asnacios, and Jean-Baptiste Manneville. Multiscale rheology of glioma cells. *Biomaterials*, 275: 120903, August 2021. ISSN 0142-9612. doi: 10.1016/j.biomaterials.2021.120903.
14. Uwe Schmidt, Martin Weigert, Coleman Broaddus, and Gene Myers. Cell detection with star-convex polygons. *Lecture Notes in Computer Science (including subseries Lecture Notes in Artificial Intelligence and Lecture Notes in Bioinformatics)*, 11071 LNCS:265–273, 2018. ISSN 16113349. doi: 10.1007/978-3-030-00934-2_30/TABLES/1. arXiv: 1806.03535 Publisher: Springer Verlag ISBN: 9783030009335.
15. Jean Yves Tinevez, Nick Perry, Johannes Schindelin, Genevieve M. Hoopes, Gregory D. Reynolds, Emmanuel Laplantine, Sebastian Y. Bednarek, Spencer L. Shorte, and Kevin W. Eliceiri. TrackMate: An open and extensible platform for single-particle tracking. *Methods*, 115:80–90, February 2017. ISSN 1046-2023. doi: 10.1016/J.YMETH.2016.09.016. Publisher: Academic Press.

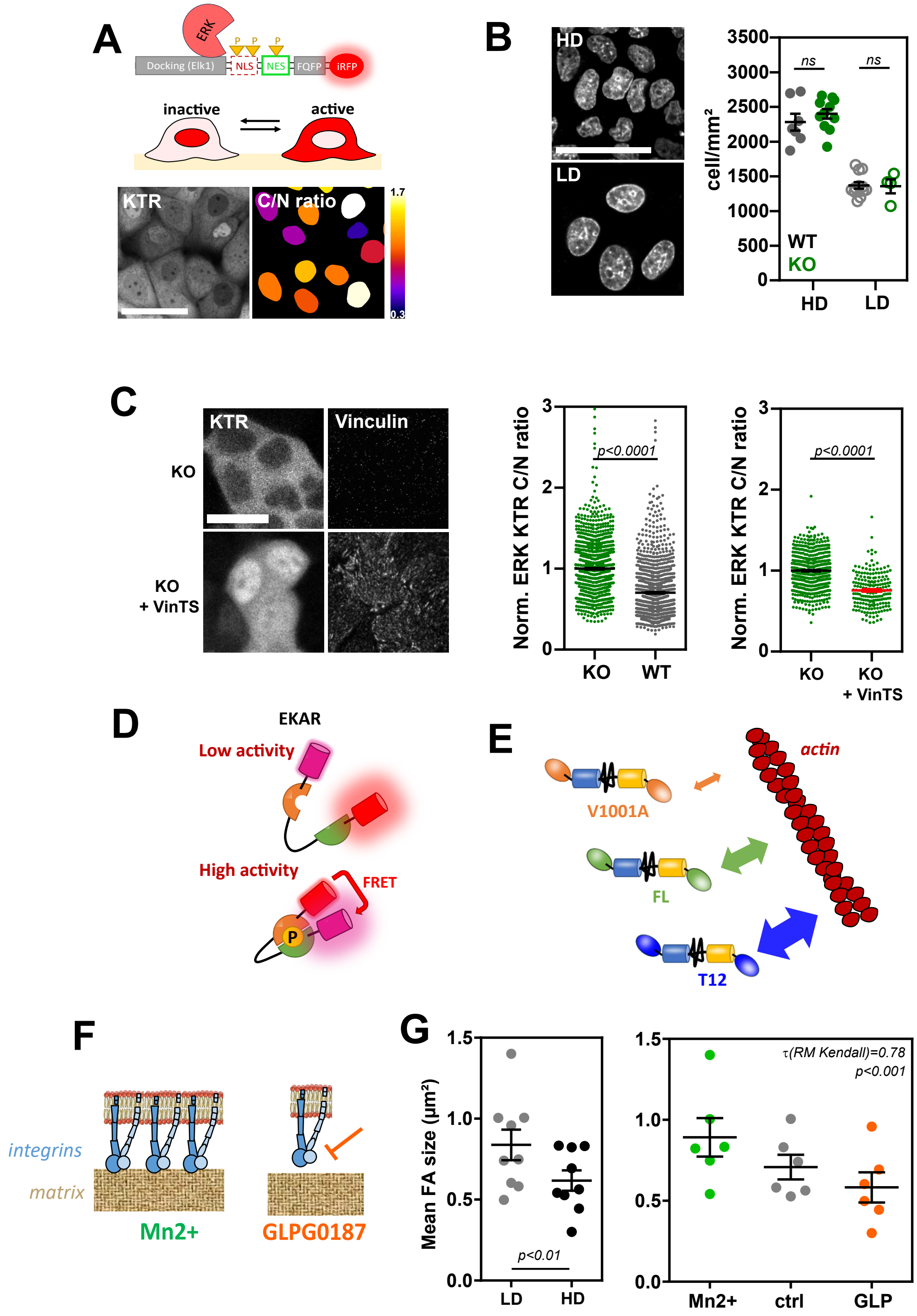


**Fig. S 1.** (A) Top: sketch of the ERK-KTR-iRFP sensor and its expected behavior. Bottom: fluorescence image of epithelial cells at confluence expressing the ERK-KTR sensor and its corresponding Cytoplasm/Nucleus intensity ratio. Scale bar=50$\mu$m. (B) Left: typical images of nucleus-stained cells at high (HD) and low (LD) epithelial densities from the same experiments as Fig. 1A. Scale bar=50$\mu$m. Right: cell densities measured from nucleus staining in HD and LD conditions for WT and KO cells. Mean$\pm$SEM of 19 (WT) and 15 (KO) FOVs from 2 (WT) and 3 (KO) independent experiments (Same as in Fig. 1A). (C) Left: fluorescence images of the ERK-KTR sensor in KO cells rescued (or not) with the VinTS construct. Scale bar=20$\mu$m. Right: normalized ERK-KTR C/N ratio in KO versus WT cells and in KO cells rescued (or not) with VinTS. Mean $\pm$SEM of $> 100$ cells per condition from 3 independent experiments. (D) Sketch of the EKAR FRET sensor for ERK activity bearing mRuby and mKate fluorophores. (E) Sketch of VinTS mutants V1001A and T12 and their expected effect on F-actin binding compared to the control VinTS (FL). (F) Sketch of the expected effects of integrin activation ($Mn^{2+}$) and inhibition (GLPG0187) treatments. (G) Mean Focal Adhesion (FA) size as a function of epithelial density (LD/HD) and integrin activation or inhibition ($Mn^{2+}$/GLP). Mean $\pm$SEM of 9 (LD, HD) and 6 ($Mn^{2+}$, ctrl, GLP) independent experiments ($> 900$ FAs per point). Two-tailed Mann-Whitney (B, C), paired Mann-Whitney (G, left), and RM Kendall (G, right) tests.

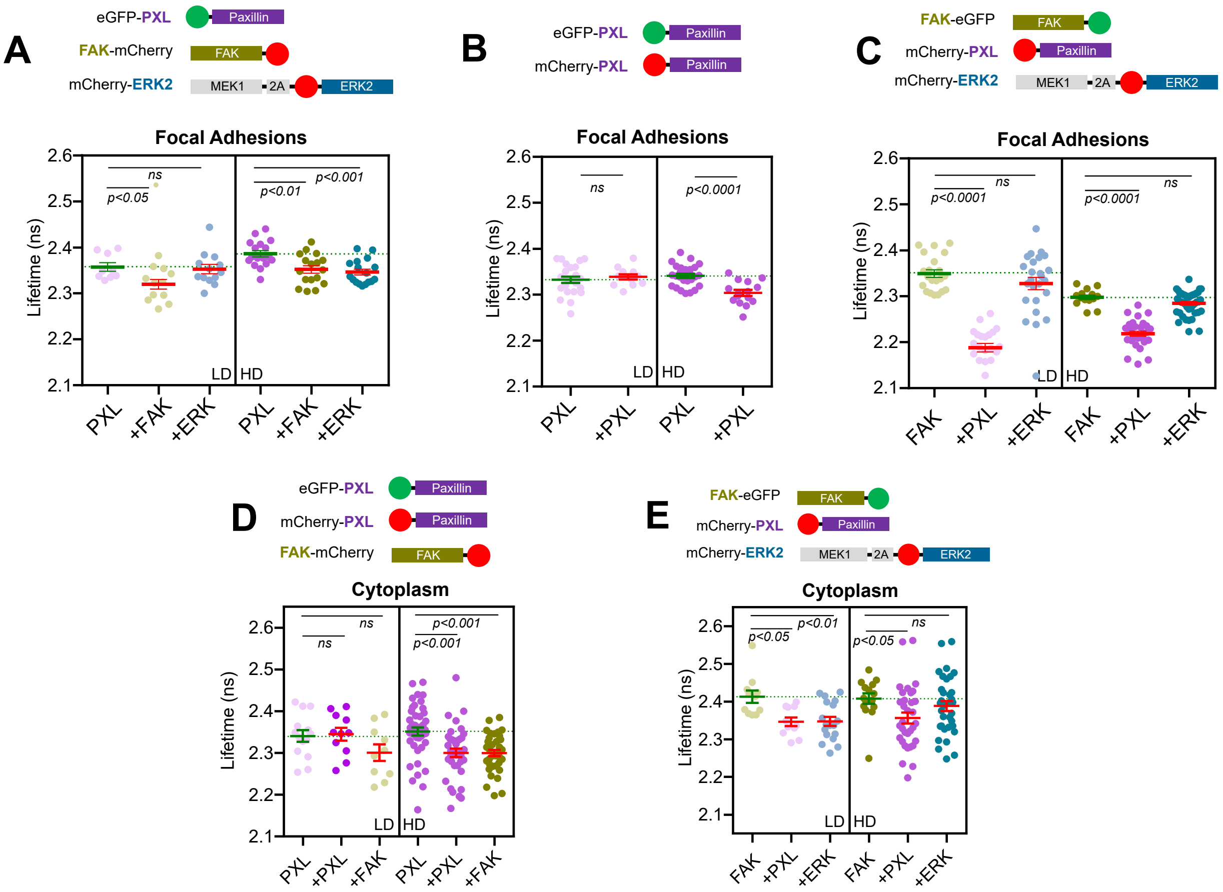


**Fig. S 2.** (A-D) Top: sketch of the constructs used for FRET-FLIM experiments. Bottom: Fluorescence lifetime of PXL-eGFP (A, B, D), FAK-eGFP (C, E) in Focal Adhesions (A-C) and in the cytoplasm (D, E) of cells expressing another mCherry construct or none at low (LD) and high (HD) cell density. Mean$\pm$SEM of 9 (A, LD, PXL), 12 (A, LD, +FAK), 13 (A, LD, +ERK), 17 (A, HD, PXL), 17 (A, HD, +FAK), 17 (A, HD, +ERK), 23 (B, LD, PXL), 12 (B, LD, +PXL), 26 (B, LD, PXL), 15 (B, HD, +PXL), 21 (C, LD, FAK), 22 (C, LD, +PXL), 25 (C, LD, +ERK), 14 (C, HD, FAK), 30 (C, HD, +PXL) and 43 (C, HD, +ERK) cells ($> 30$ FAs) from 2 independent experiments, and 14 (D, LD, PXL), 11 (D, LD, +PXL), 10 (D, LD, +FAK), 44 (D, HD, PXL), 38 (D, HD, +PXL), 36 (D, HD, +FAK), 11 (E, LD, FAK), 11 (E, LD, +PXL), 18 (E, LD, +ERK), 15 (E, HD, PXL), 33 (E, HD, +PXL) and 34 (E, HD, +FAK) cells from 2 independent experiments. Kruskal-Wallis and Dunn's multiple comparisons (A, C-E), and Mann-Whitney (B) tests.

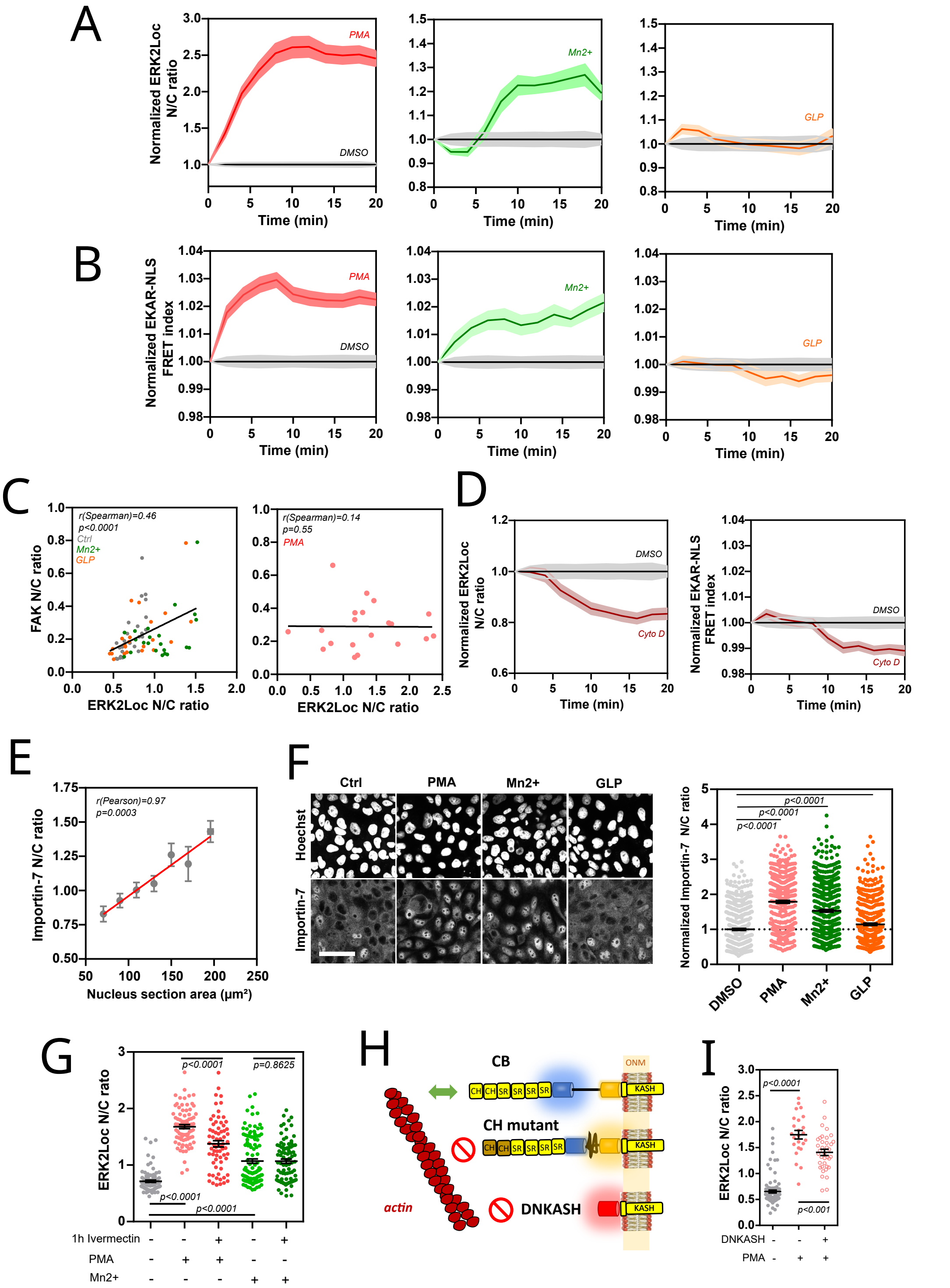


**Fig. S 3.** (A, B) Kinetics of ERK nuclear translocation (ERK2Loc N/C ratio) and activity (EKAR-NLS FRET index) upon pharmacological activation of ERK (PMA) or upon activation ($Mn^{2+}$) or inhibition (GLP) of integrins. Mean ±SEM of 46 (DMSO), 26 (PMA), 33 ($Mn^{2+}$) and 25 (GLP) cells from 3 independent experiments (A), and 65 (DMSO), 42 (PMA), 14 ($Mn^{2+}$) and 46 (GLP) cells from 2 ($Mn^{2+}$) or 3 independent experiments (B). (C) Correlation between FAK and ERK nuclear translocation, spontaneous and upon activation ($Mn^{2+}$) or inhibition (GLP) of integrins (left), or upon pharmacological activation of ERK (PMA). 25 (Ctrl), 22 ($Mn^{2+}$), 22 (GLP) and 20 (PMA) cells from 2 independent experiments. (D) Kinetics of ERK nuclear translocation (ERK2Loc N/C ratio) and activity (EKAR-NLS FRET index) upon cytoskeleton disruption (CytoD). Mean ±SEM of 46 (DMSO) and 45 (CytoD) from 3 independent experiments ERK2Loc), and 65 (DMSO) and 48 (CytoD) from 3 independent experiments (EKAR-NLS), same DMSO reference as in (A, B). (E) Importin-7 immunofluorecence intensity N/C ratio as a function of nucleus section area. Mean ±SEM of 62, 93, 100, 109, 66, 24 and 11 cells per 20$\mu m^2$-bin between 70 and 190 $\mu m^2$ from 2 independent experiments. (F) Left: importin-7 immunofluorescence images as function of pharmacological ERK activation (PMA) or of integrin activation ($Mn^{2+}$) or inhibition (GLP). Right: Importin-7 N/C ratio in these conditions. Mean ±SEM of 497 (Ctrl), 462 (PMA), 444 ($Mn^{2+}$) and 403 (GLP) cells from 2 independent experiments. (G) ERK2Loc N/C ratio as a function of importin-$\alpha/\beta$ inhibition (Ivermectin), ERK activation (PMA) and integrin activation ($Mn^{2+}$). Mean ±SEM of 90 (Ctrl), 81 (PMA), 68 (Ivermectin+PMA), $Mn^{2+}$ (116) and 88 (Ivermectin+$Mn^{2+}$) cells from 2 independent experiments. (H) Sketch of the nesprin constructs: cytoskeleton-binding (CB) mN2G-TS, calponin-homology domain mutant mN2G-TS (CH), and dominant negative DNKASH and their expected interaction with actin. (I) ERK2Loc N/C ratio as a function of LINC complex disruption (DNKASH) and pharmacological activation of ERK (PMA). Mean ±SEM of 99 (Ctrl), 24 (PMA) and 37 (DNKASH+PMA) cells from 2 independent experiments. Two-tailed Mann-Whitney tests (F, G, I) and Spearman (C) or Pearson correlations (E).

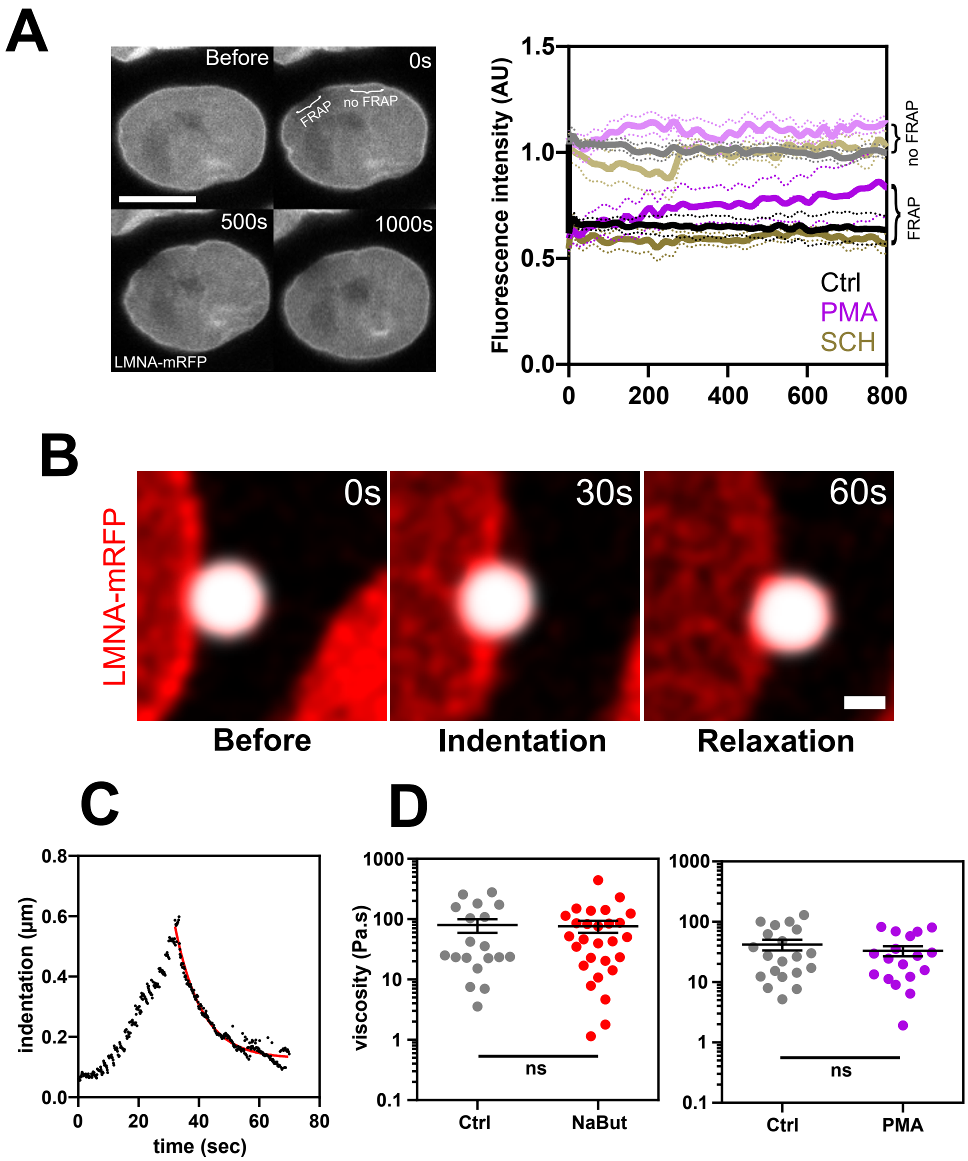


**Fig. S 4.** (A) Left: images of a LMNA-mRFP nucleus before and during FRAP. Bar$= 10\mu$m. Right: fluorescence intensity of LMNA-mRFP in FRAPPED and non-FRAPPED regions of the nuclear envelope in ERK-activated (PMA) or -inhibited (SCH) cells compared to control. Mean $\pm$SEM of 17 (Ctrl), 8 (PMA) and 11 (SCH) cells from 3 (Ctrl, PMA) and 2 (SCH) independent experiments. (B) Images of a LMNA-mRFP nucleus (red) in a live cell before, during and after indentation with a $2\mu$m bead (gray) trapped with the optical tweezer. Bar$= 1\mu$m. (C) Typical variation of the nucleus indentation during an indentation-relaxation sequence. The red line shows the exponential decay fit to extract the viscosity from the relaxation phase. (D) Effects of deacetylase inhibition with Sodium Butyrate (NaBut) and ERK activation (PMA) on viscosity. Mean $\pm$SEM of 19 (Ctrl/NaBut), 28 (NaBut), 20 (Ctrl/PMA) and 18 (PMA) from 4 independent experiments. Two-tailed Mann-Whitney tests.

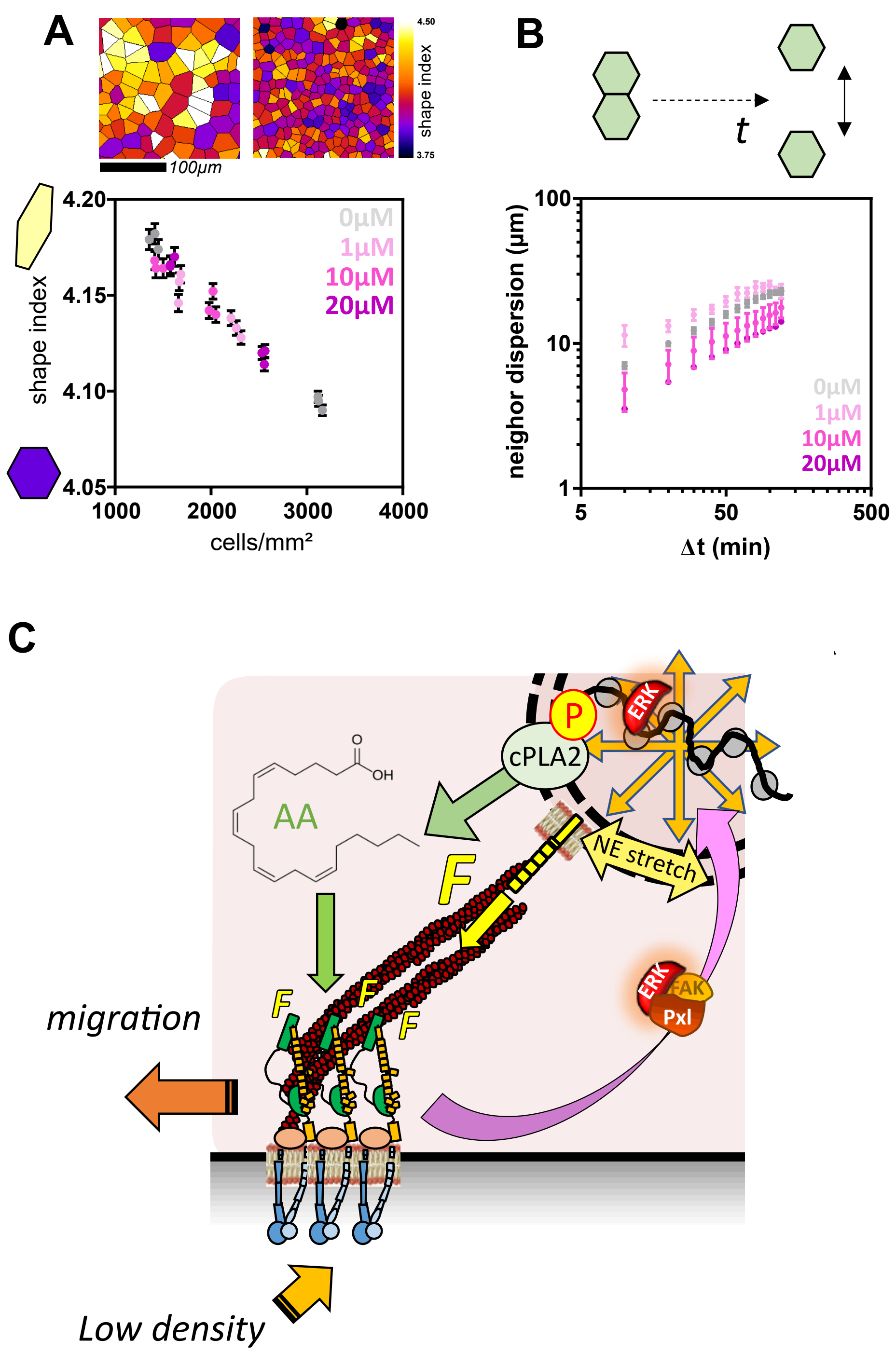


**Fig. S 5.** (A) Top: Voronoi tesselation from nuclei centers in cells at confluence at about 1500 and 3000 cells/mm$^2$ and their respective shape indices. Bottom: shape index as a function of cell density and cPLA2 inhibition. Mean $\pm$SEM of $> 1000$ cells per FOV from 3 (Ctrl, 1$\mu$M) and 2 (10, 20 $\mu$M) experiments (same as in Fig. 5D). (B) Top: Sketch of the neighbor dispersion metric. Bottom: Neighbor dispersion for increasing levels of cPLA2 inhibition (AACOCF3 concentration). Mean $\pm$SEM of 3 (1$\mu$M) and 2 (Ctrl, 10, 20 $\mu$M) FOVs ($> 100$ cells), each from an independent experiment (same as in Fig. 5D). (C): Working model of the control of cell migration by epithelial density through a mechanotransduction pathway involving cPLA2 activation upon ERK translocation into the nucleus. See text for details.
